# High-throughput field phenotyping reveals that selection in breeding has affected the phenology and temperature response of wheat in the stem elongation phase

**DOI:** 10.1101/2022.09.05.506627

**Authors:** Lukas Roth, Lukas Kronenberg, Helge Aasen, Achim Walter, Jens Hartung, Fred van Eeuwijk, Hans-Peter Piepho, Andreas Hund

**Affiliations:** ETH Zurich, Institute of Agricultural Sciences, Universitätstrasse 2, 8092 Zurich, Switzerland; Agroscope, Earth Observation of Agroecosystems Team, Division Agroecology and Environment, Reckenholzstrasse 191, 8046 Zurich, Switzerland; University of Hohenheim, Institute for Crop Science, Biostatistics Unit, Fruwirthstrasse 28, 70593 Stuttgart, Germany; Wageningen University and Research, Biometris, P.O. Box 16, 6700 AA Wageningen, The Netherlands

**Keywords:** Correlated response to selection, Genetic correlation, Genomic prediction, Growth dynamic, GWAS, Modeling, Trait extraction

## Abstract

Crop breeders increasingly need to mitigate the effects of climate change. Ideally, their selection strategies are based on an understanding of crop responses to environmental covariates such as temperature. In this study, the height of 352 varieties (European and Swiss) was repeatedly measured in multiple years. P-splines were used to model plant height as a function of time, from which the phenology parameters jointing (start) and end of stem elongation were derived. An asymptotic model was used to estimate the base-temperature of growth (*T*_min_), the steepness of the response (*lrc*), and the growth at optimum temperature (*r*_max_). Parameter *r*_max_ had the largest effect on final height, whereas the temperature-response parameters in the narrow sense (*T*_min_, *lrc*) were closely connected to phenology. Final height and *r*_max_ decreased from the more continental, eastern European countries towards the more maritime, western countries. For genotypes registered in Great Bitain, *T*_min_ was distinctly lower compared to most other regions. Integrating such analysis of in-season responsiveness to fluctuating environmental conditions in breeding will help to improve the genetic gain for climate adaptation. It can only be achieved based on high-throughput assessment of phenotypes in the field throughout the season.

**Highlight:** Wheat phenology and environmental responses have been affected by breeders’ selections since 1970. High throughput field phenotyping methods reveal these developments, allowing to better adapt future varieties to climate change.

## 1. Introduction

Mitigating climate change impacts on crops through genotypic adaptation requires understanding crop responses to environmental factors (Ramirez-Villegas et al., 2015). Responses of major crops are well studied in controlled environments but the translation of insights to the field is not straightforward (Poorter et al., 2016). High-throughput field phenotyping (HTFP) may facilitate this transition (Araus et al., 2018).

A main driver of plant growth and development is temperature (Porter and Gawith, 1999). By examining 17 crop species including wheat (*Triticum aestivum* L.), Parent and Tardieu (2012) have not found a relation between breeding and temperature responses. In contrast, Kronenberg et al. (2020a) found genotype-specific temperature responses for a set of European winter wheat genotypes in the field (the GABI-Wheat panel, Kollers et al. 2013; Gogna et al. 2022). A crucial difference between the two investigations is that the former normalized growth rates of genotypes to unity at 20 °C while the latter did not. Thus, Kronenberg et al. (2020a) have used an approach that allowed genotypes to differ in growth rates at optimal temperature.

Gaining insights on adverse aspects of temperature response and phenology (i.e., the timing of key stages) is of high interest for breeding. Increasing the duration of stem elongation by adjusting heading time or the beginning of stem elongation (jointing) has repeatedly been proposed as possibility to increase wheat yield (Slafer et al., 1996; Miralles and Slafer, 2007). Phenology is driven by environmental and genotype characteristics and corresponding interactions, and therefore a result of G×E. In contrast, temperature response traits are only to a limited extent affected by—but rather drivers of—G×E. Describing such responses directly in the breeding nursery may allow breeders to predict the phenotypic performance in new unseen environments (Poorter et al., 2010). Yet, differences between development of wheat varieties originating from various world regions are not well documented and understood (Gbegbelegbe et al., 2017). CIMMYT has defined 17 ‘Mega-Environments’ for wheat production (Monfreda et al., 2008) and only recently, it was shown that using region-specific cultivar parameters is critical when applying crop models at a global scale because cultivars vary in response to climate conditions, soils and crop management (Gbegbelegbe et al., 2017). The extent to which genotypes differ is unclear because it is difficult to trace the development of wheat at a relevant scale. With the advent of high-throughput phenotyping methods, this has become feasible nowadays. One of the most simple traits to detect is plant height. This trait can be analyzed at a temporal resolution of a few days with several methods. Kronenberg et al. (2020a) used a terrestrial laser scanner mounted on a rope-suspended phenotyping platform (Kirchgessner et al., 2017) to determine plant height. From a breeder’s perspective, such a stationary platform is highly inflexible as it does not allow screening multi-environment trials. Mobile platforms are better suited to screen large breeding populations (Aasen and Bareth, 2018), allowing increasing the genetic gain of selection (Araus et al., 2018). Thus, the first aim of this study was to test whether a drone-based phenotyping platform, carrying RGB-cameras (Roth et al., 2018) allows extracting temperature response traits from the GABI-Wheat panel with comparable repeatability as achieved by laser scanning.

Independent of the measurement device, modeling the temperature response from field-derived data bears flaws and pitfalls (Roth et al., 2022b). The appropriateness of a dose-response curve does not only depend on the (biological) response but also on the range of measured temperatures. Using a linear regression to model temperature response as done in Kronenberg et al. (2020a) is controversial: Such a Type 1 response (Wang et al., 2017) will come to its limits when measurements span a whole growing season with temperatures also extending into supra-optimal ranges (Parent et al., 2019; Kronenberg et al., 2020a). Using an asymptotic temperature response curve may overcome this limitation and allows resolving the temperature response into a base temperature (*T*_min_), the growth rate at optimal temperatures (*r*_max_), and the steepness of the temperature response (*lrc*) (Roth et al., 2022b). An aim of this study was to evaluate whether the asymptotic model is suitable to extract meaningful parameters from real high-throughput field phenotyping (HTFP) data.

The GABI-Wheat panel (Kollers et al., 2013; Gogna et al., 2022) includes important genotypes from different climatic regions of Europe, having all been registered in Mega-Environment 11 of the CIMMYT convention. Since European countries have pursued country-specific, largely independent breeding programs, it is interesting to look at the development of phenology in a large population of such cultivars during recent decades that have passed under the advent of global climate change. The adjustment of phenology is a major breeding aim which leads to a strong population structure at genetic loci related to phenology. This raises the question: Which traits were the target of selection, and which traits only show a correlated response? Genetic correlations can be used to analyze the correlated response of traits to selection (Falconer and Mackay, 1996). In contrast, grouping phenotypes by variety registration country and year may allow examining population structures. Finally, genome wide association studies (GWAS) allow gaining insight into the genetic architecture of the investigated traits as well as their interrelations.

Hence, an additional aim of this study was to characterize the GABI-Wheat panel using phenotypic and genetic correlations and GWAS, providing insights on direct and indirect response to selection in breeding programs but also on general genetic relationships.

## 2. Materials and Methods

### 2.1. Experimental design

Experiments were performed in four consecutive years (2015 to 2018) in the field phenotyping platform FIP (Kirchgessner et al., 2017) at the ETH research station of agricultural sciences in Lindau Eschikon, Switzerland (47.449 N, 8.682 E, 556 m a.s.l.). Details about designs, genotypes, soil and management can be found in Kronenberg et al. (2017, 2020a) for 2015 to 2017 and Roth et al. (2020) for 2018.

In brief, a GABI-Wheat subset (consisting of 300 European winter wheat cultivars from the GABI-Wheat panel, Kollers et al. (2013); Gogna et al. (2022)) was complemented by 35 to 52 Swiss winter wheat varieties of commercial importance. The resulting panel of in average 345 genotypes was replicated twice per year and each replication was planted on a different lot in the FIP area. Each replication was augmented with spatial checks in a 3×3 block arrangement (Figure 1). Designs were enriched with spatial coordinates based on unmanned aerial system (UAS) flights for the years 2017 and 2018. For 2015 and 2016, no UAS flights were available and therefore local coordinates (row and range) were used as spatial context as described by Kronenberg et al. (2020a).

**Figure 1:**
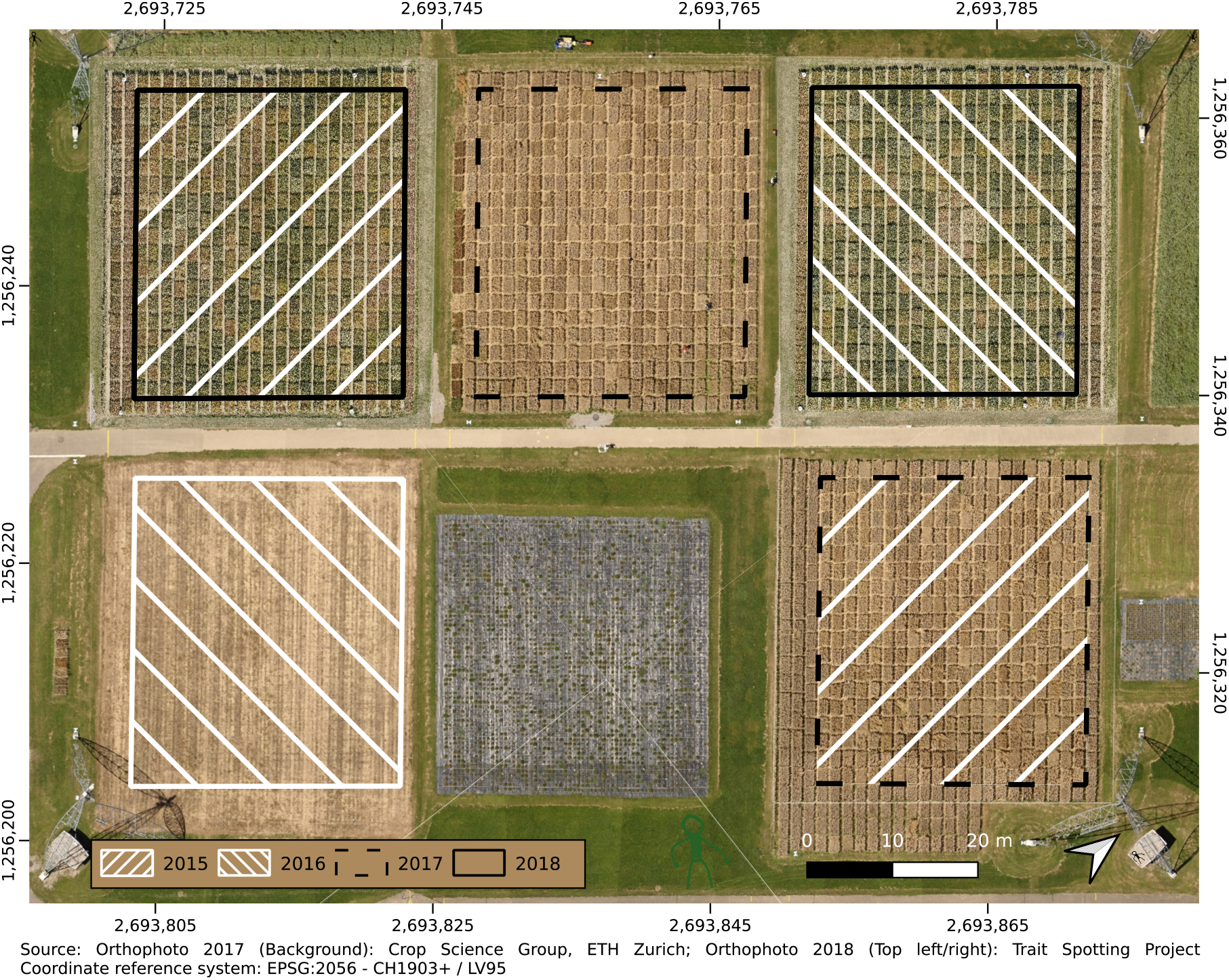
Experimental fields at the FIP site with indications of the two replications per year (boxes) and aerial images of two years (2017, superimposed on 2018 top left and top right) as background.

### 2.2. Phenotyping and covariate measurements

Plant height measurements for the years 2015 to 2017 were performed using a terrestrial laser scanner (TLS) based on a light detection and ranging (LiDAR) sensor (FARO R Focus3D S 120; Faro 113 Technologies Inc., Lake Mary, U.S.A.) mounted on the FIP (Kronenberg et al., 2017). Measurements were performed every three to four days from jointing to harvest in 2015 and 2016 and from tiller development to harvest in 2017 (Kronenberg et al., 2020a).

For the year 2018, FIP measurements were replaced by the UAS platform PhenoFly (Roth et al., 2018). The UAS captured RGB images with high spatial overlap that were processed using Structure-from-Motion (SfM) software (Agisoft Metashape, Agisoft LLC, St. Petersburg, Russia) to yield digital plant height models. General flight campaign settings are described in Roth et al. (2020). Flights were performed every two to three days from tiller development to harvest.

To ensure comparability between TLS and UAS data, measurements were performed with both platforms simultaneously on five dates in 2018 (04-06, 04-11, 04-19, 05-09, 06-14) for one replication. Meteorological data were obtained from a weather station next to the experimental field (50 m). Data gaps were filled with data from a close-by public Agrometeo weather station (http://www.agrometeo.ch/, Agroscope, Nyon, Switzerland) in proximity (550 m).

### 2.3. Plant height extraction

For TLS measurements, individual plot-based plant height values were extracted from point clouds using a custom-developed Matlab script as described in Friedli et al. (2016) and Kronenberg et al. (2017). For UAS measurements, plot-based values were extracted in Python as described by Roth and Streit (2018) with one modification: Before processing digital elevation models (DEMs) to plant height models, DEMs were spatially corrected using reference ground control point (GCP) coordinates. To do so, differences between DEM elevations and known reference elevations were calculated at all GCP locations and a cubic interpolation performed on the whole experimental area. Interpolated differences were then subtracted from the original DEM to produce a corrected DEM.

### 2.4. Dynamic modeling

#### 2.4.1. Timing of jointing and end of stem elongation

In a first step, a shape constrained monotonically increasing P-spline was fitted to plot time series using the R package *scam* (Pya, 2019) to extract the timing of jointing and the end of stem elongation (Figure 2a). The package fits shape-constrained generalized additive models (GAM) using a Bayesian framework. Standard errors of predictions are computed from the posterior distribution of fitted spline coefficients. The number of knots was fixed to 3/4 of the number of observations.

**Figure 2:**
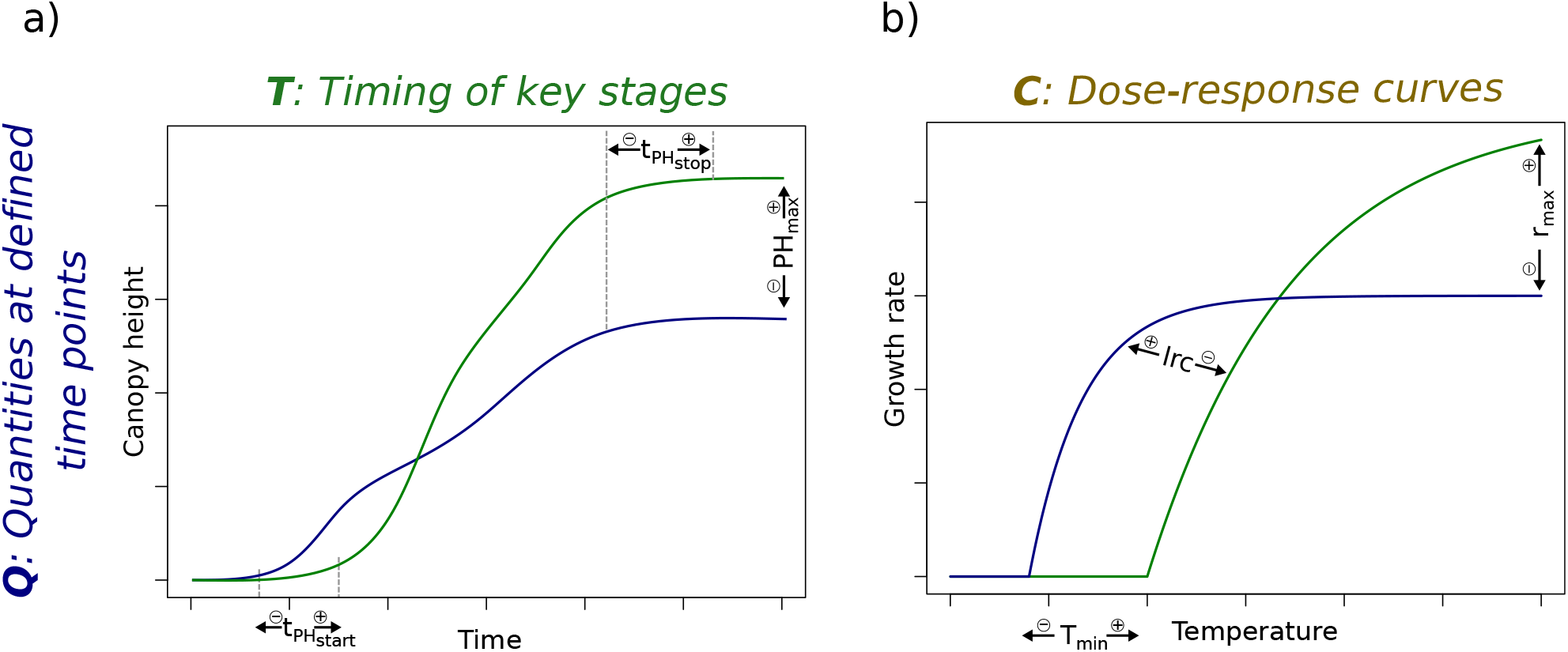
Visualization of traits derived from the spline models followed by QMER extraction of timing of key stage traits jointing (*t*_PH_start__) and end of stem elongation (*t*_PH_stop__) and quantity trait final height (PH_max_) (a) and from the asymptotic model to determine the temperature-response parameters maximum elongation rate (*r*_max_), base temperature where the elongation rate is zero (*T*_min_) and steepness of the response (*lrc*) (b).

In a subsequent step, spline predictions for canopy heights and standard errors of these estimates were calculated on an hourly timescale using the prediction function of the *scam* package. Then, the quarter of maximum elongation rate (QMER) method (Roth et al., 2021) was applied to extract the growth stages jointing (*t*_PH_start__) and end of stem elongation (*t*_PH_stop__). In brief, the QMER method determines the time point at which the elongation rate exceeds (*t*_PH_start__) or falls short of (*t*_PH_stop__) by a certain threshold of the maximum elongation rate.

#### 2.4.2. Final height

The final height (PH_max_) (according to Roth et al. (2021) a quantity at a defined time point trait) was calculated as the median of the top 24 spline predictions after the estimated stop of growth *t*_PH_stop__ (Figure 2a). For simplicity, the corresponding standard error estimation from *scam* was used for weighting in further processing, which is based on the uncertainty from the spline fitting but neglects the uncertainty of the extraction method.

#### 2.4.3. Temperature dose-response parameters

Measuring plant height with high-throughput devices allows deriving growth rates from successive measurements (Kronenberg et al., 2020a). To extract the temperature response of growth from these time series, an asymptotic model *r*_asym_ (Figure 2b) that defines growth rates *r* as a function of temperature *T* was applied to plot time series height data between *t*_PH_start__ and *t*_PH_stop__,

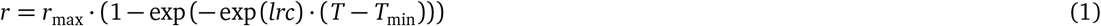

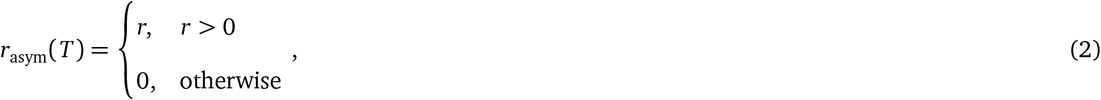

where *T* are hourly temperature means, *r*_max_ is the maximum elongation rate (and therefore the asymptote of the curve), *T*_min_ the base temperature where the elongation rate is zero, and *lrc* characterizes the steepness of the response.

Maximum-likelihood-fitting was used to fit the asymptotic dose-response curve (Equation 1, 2) to plant height measurements with irregular measurement intervals (2–4 days) and hourly temperature measurements (Roth et al., 2022b). Parameters were estimated in R using the method L-BFGS-B of the function *optim*. Note that the residuals were assumed to be identically and independently normally distributed, as previous attempts to include an autocorrelation term resulted in failed convergence. Standard errors as weightings for further processing were extracted based on the Hessian matrix provided by *optim*. Subsequently, weights for further processing were computed as inverse squared standard errors.

To allow for a comparison with previous field-based studies (Grieder et al., 2015; Kronenberg et al., 2020a), an additional linear model was fitted to the data. This model regressed growth rates on mean air temperatures of the corresponding measurement period (i.e., the mean of a 3–5 day hourly temperature time series), thus extracting the temperature response parameter lm_slope_ which corresponds to the slope reported in Kronenberg et al. (2020a).

### 2.5. Adjusted genotype means per year and repeatability

The above described dynamic modeling of plot-based repeated measures into plot-based intermediate traits can be seen as a first stage of stage-wise processing (Roth et al., 2021). These plot-based intermediate traits were further processed in a stage-wise linear mixed model analysis (stage two and three), in which the second stage averaged over within-year effects (resulting in adjusted genotype-year means) and the third stage over between-year effects (resulting in overall adjusted genotype means). For the second stage, the R package SpATS (Rodríguez-Álvarez et al., 2018) was parameterized with the model

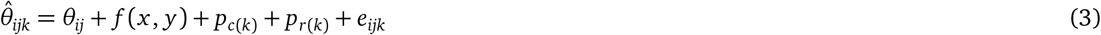

for each year *j* separately, where 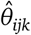 are plot responses based on dynamic modeling for the *i*th genotype, *j*th year, and *k*th replication, *θ_ij_* genotype-year responses, *p*_*c*(*k*)_ random effects in range direction per replication *k* (main working direction, e.g. for sowing), *p*_*r*(*k*)_ random effects in row direction per replication *k* (orthogonal to main working direction), *f* (*x,y*) a smooth bivariate surface in spatial *x* and *y* coordinates consisting of a bivariate polynomial and a smooth part (for details see Rodríguez-Álvarez et al., 2018), and *e_ijk_* plot residuals assumed to have variance equal to the squared standard error estimations from the previous dynamic modeling. Weighting was only applied when decreasing the Bayesian information criterion (BIC) if compared to a model without weighting. Weights did not improve BIC for 2018 for *r*_max_, for all years for *T*_min_, for 2015–2017 for *lrc*, for 2017 for *t*_PH_start__, and for 2015, 2016 and 2017 for *t*_PH_stop__.

For best linear unbiased estimation (BLUE) calculations, *θ_ij_* was set as fixed, all other terms as random. For best linear unbiased prediction (BLUP) and repeatability calculations, *θ_ij_* was set as random. Within-year heritability (repeatability) calculation based on BLUPs was calculated according to Oakey et al. (2006).

### 2.6. Across-year adjusted genotype means and heritability

For the third stage, the R package ASReml-R (Butler, 2018) was parametrized with the model

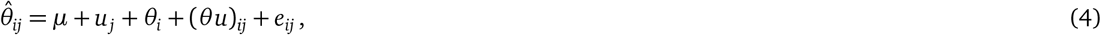

where 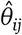 are adjusted year genotype means (BLUE) from the previous stage, *μ* a global intercept, *u_j_* year effects, *θ_i_* genotype responses, (*θu*)_*ij*_ genotype year interactions allowing for year specific variances (diagonal variance structure), and *e_ij_* residuals weighted based on the inverse of the diagonal of the variance-covariance matrix from the previous stage.

For BLUEs calculations, *μ* and *θ_i_* were set as fixed, all other terms as random. For BLUPs and heritability calculations, *θ_i_* was set as random with known variance structure based on the normalized genome-wide average identity by state (IBS) relationship structure calculated from single nucleotide polymorphism (SNP) marker data using the *snpgdsIBS* function in the R-package SNPRelate (Zheng et al., 2012).

Marker data was supplied by the GABI wheat consortium (Kollers et al., 2013; Gogna et al., 2022) for the GABI wheat genotypes and by Agroscope in the framework of the Swiss winter wheat breeding program (Fossati and Brabant, 2003) for the Swiss genotypes. For the IBS analysis, only non-monomorphic SNPs with unequivocal genome positions (see Kronenberg et al., 2020a) and a missing rate < 0.05 and a minor allele frequency < 0.05 were used, thus resulting in 9147 SNPs for 325 genotypes. Heritability was calculated on a genotype-difference basis following the 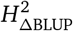 method defined in Schmidt et al. (2019).

### 2.7. Phenotypic and genetic correlation calculation

The phenotypic correlations between traits were calculated for each of the four examined years as Pearson’s r of plot-based values. For reporting, the mean, maximum and minimum of these four correlations per trait pair were calculated. For the genetic correlation calculation, the univariate model of Equation 4 was extended to a bivariate model (Wright, 1998; Holland et al., 2001),

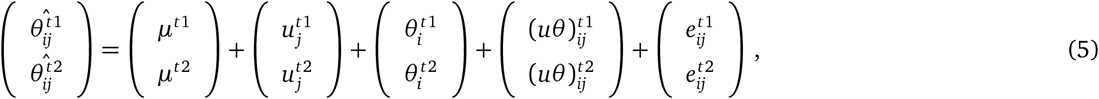

where 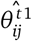 and 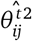 are adjusted year genotype means (BLUEs) per trait (trait 1 (*t*1) and trait 2 (*t*2)) from the second stage, *μ*^*t*1^ and *μ*^*t*2^ global intercepts per trait, 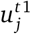 and 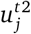 year effects per trait, 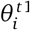 and 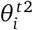 genotype responses (with known variance structure based on IBS), and (*uθ*^*t*1^)_*ij*_ and (*uθ*^*t*2^)_*ij*_ the genotype responses to 3 year interactions with uniform variances per trait. The terms 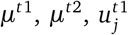 and 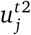 were set as fixed, all other terms were random. Note that no variance-covariance-matrix from the previous stage for a bivariate model was available. Consequently, *e* and (*uθ*) are confounded, whence the two terms were summarized in one variance-covariance structure. Genetic correlations among traits were then calculated based on the estimated variance and covariance components (Holland et al., 2001),

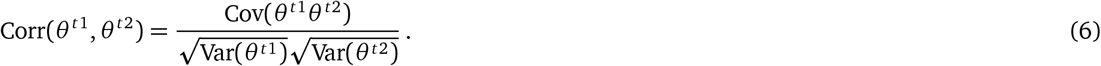

To investigate the effect of release year and country of origin on intermediate traits, the year and country of first registration of genotypes was looked up in the EU plant variety database (https://ec.europa.eu/food/plant/plant_propagation_material/plant_variety_catalogues_databases). Five multi-year-groups ((1970-1990], (1990,1995], (1995, 2000], (2000,2005], and (2005,2018]) and seven countries groups (AT/CZ, PL, DE, CH, SE/DK, FR, GB) were chosen and phenotypic values per group aggregated to means and standard errors. Note that only country-groups with a sample size ≥10 were considered, and that the first and last year clusters have, due to the focus of the GABI Wheat panel on the release years 1990–2005, wider ranges than the other clusters.

### 2.8. Genomic prediction and genome wide association studies

In a next step, the suitability of the extracted intermediate traits for genomic prediction was estimated. To align results with existing literature (e.g., Bustos-Korts et al., 2019; Meher et al., 2022; Toda et al., 2022), overall genotype means (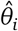 in Equation 4) were used as phenotypic values.

The prediction accuracy was evaluated based on a Genomic Unbiased Prediction (GBLUP) model parametrized in the R package ASReml-R (Butler, 2018),

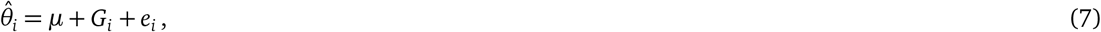

where 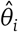 are across-year BLUEs (Equation 4), *μ* a global intercept, and *G_i_* are random genotype effects with *G* = (*G*_1_, *G*_2_,…)^*T*^ based on a variance-covariance-matrix calculated as the IBS relationship structure mentioned in the previous section (Section 2.6). Residuals *e_i_* were weighted based on the inverse of the diagonal of the variance-covariance matrix from the previous stage (Equation 4). Prediction accuracy was calculated as mean Pearson’s r of ten-fold cross validations, random folds were ten times repeated.

To investigate the underlying genetic architecture of the different traits and assess the observed phenotypic and genetic correlations in this context, we performed genome wide association studies (GWAS) on the 325 wheat varieties present across all year-sites. The same genotype data was used as for the IBS analysis (see Section 2.6). Univariate GWAS were conducted for all traits using three different models implemented in the R package *GAPIT* (Wang and Zhang, 2021). As a baseline approach, a single locus, compressed, mixed linear model (*MLM*) (Zhang et al., 2010) was used including the first three principal components of the marker genotypes as fixed effects and a kinship matrix calculated following VanRaden (2008) as random effects. Further, the two multi-locus models *FarmCPU* (Liu et al., 2012) and *Blink* (Huang et al., 2018) were applied. While *MLM* effectively controls type I errors, the incorporated kinship and population structure can reduce the detection of true associations, especially for complex traits associated with population structure (Liu et al., 2012; Atwell et al., 2010). *FarmCPU* and *Blink* have increased power as they reduce this confounding while maintaining the control of type I errors of *MLM* (Liu et al., 2012; Huang et al., 2018).

In order to investigate putative pleiotropic structures among the observed traits and account for the correlation structure between traits, we conducted multivariate GWAS using the software *GEMMA* (Zhou and Stephens, 2014). To this end, traits (excluding lm_slope_) were grouped into two physiological and correlation-based multitrait combinations, i) temperature response traits (*r*_max_,*T*_min_, *lrc*) and final height; and ii) phenology traits (*t*_PH_start__ and *t*_PH_stop__) and final height. For all multivariate GWAS, the first three principal components were included to correct for population structure and the same IBS matrix as used for genetic correlations was applied to correct for relatedness.

All GWAS were conducted on adjusted genotype means per year-site (BLUEs), as well as for across-year adjusted genotype means (BLUEs and BLUPs). For the detection of significant marker-trait-associations (MTAs), a Bonferroni threshold (*α* = 0.05, -log10(*P*) = 5.26) was applied to stringently correct for multiple testing.

## 3. Results

### 3.1. Plant height measurements reveal characteristics of growth dynamics

A total of 72,278 plant height estimation data points were extracted from TLS point clouds and UAS based digital elevation models, corresponding to 2,936 plot-based time series (Figure 3). These plant height time series exhibited a strong increase after the start phase in the early season, and a clear plateau after reaching the maximum height mid-season. In 2016, time series indicated lodging for specific plots at the end of this extraordinary wet growing season. The start phase of stem elongation exhibited a clear lag in the second half of April 2017, but not for other years. The dynamics of the end phase of stem elongation visually did not differ between years. Final heights clearly differed between years with tall plants in the wet year 2016 and short plants in the extraordinary dry year 2018.

**Figure 3:**
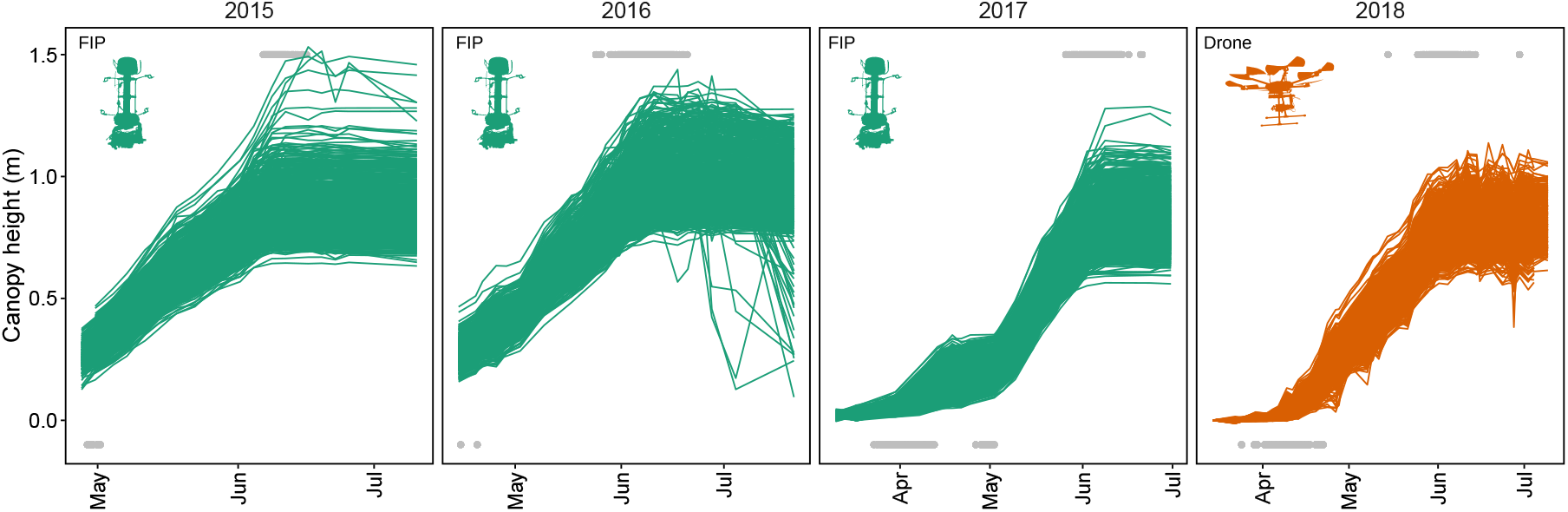
Plant height measurements performed with the field phenotyping platform (FIP, green) using a terrestrial laser scanner and with the PhenoFly platform (UAS, orange) using Structure-from-Motion based on RGB images. Grey points indicate detected start (bottom row) and end (top row) points of growth phases (*t*_PH_start__ and *t*_PH_stop__) (Note that *t*_PH_start__ for 2015 and 2016 was not reliably detected, thus 2015 and 2016 *t*_PH_start__ values were skipped for subsequent analyses).

TLS and UAS measurements performed in parallel in 2018 revealed good correlations with moderate coefficients of determination for three early dates (R^2^= 0.5-0.7) and strong coefficients of determination for two later dates (R^2^= 0.87-0.89) (Figure 4). Intercepts of the first two dates were close to zero, negative for the two subsequent dates (−0.01, −0.04 m) and positive for the last date (0.075 m), slopes indicated a severe underestimation of height by UAS measurements for early dates but weaker underestimation for later dates.

**Figure 4:**
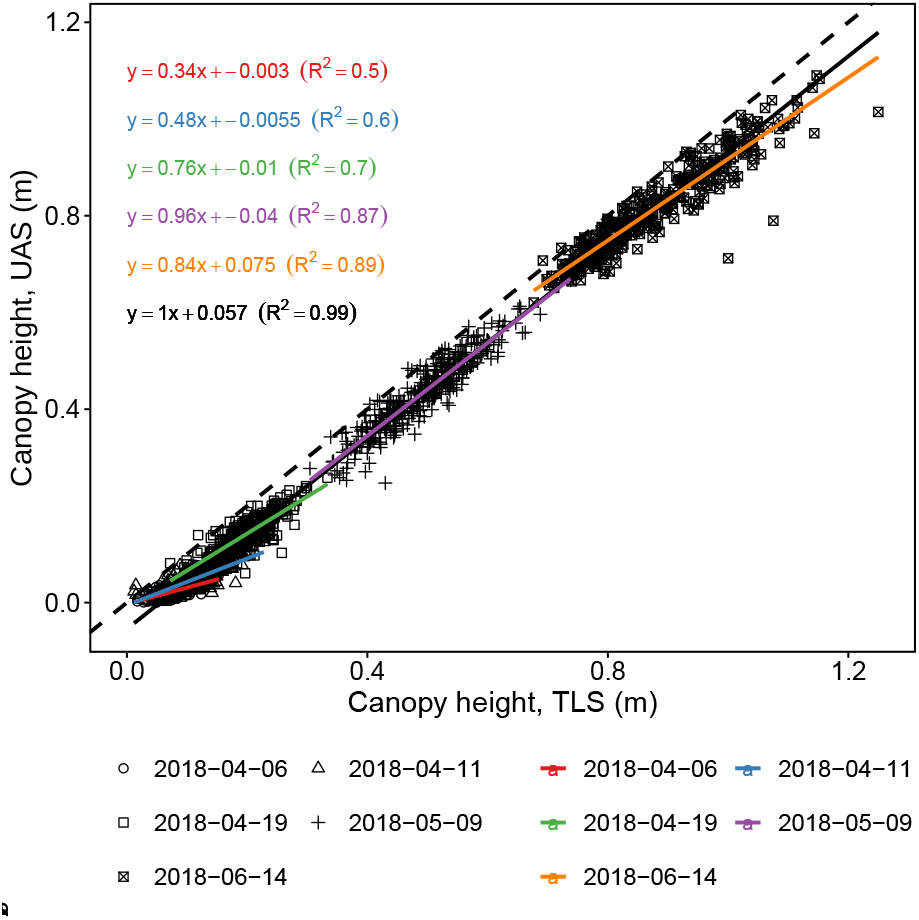
Comparison of terrestrial laser scanning (TLS) based plant heights and drone (UAS) based plant heights for five measurement dates in 2018. Colored lines are linear regressions for corresponding dates, the black line a linear regression for all dates, the colored and black text lines denote slope, intercept and goodness of fit (*R*^2^) for the corresponding linear regressions, the dashed line annotates a 1:1 relationship.

### 3.2. Dynamic modeling allows extracting heritable timing of jointing, end of stem elongation and final height traits

Fitted P-splines indicated a clear plateau after reaching final height (*t*_PH_stop__/PH_max_) (Supplementary materials, Figure S1b). Visualizing the first derivative of the splines revealed a non-steady growth phase with severe changes in growth rates (Supplementary materials, Figure S1a). Applying the QMER method to determine the timing of jointing and end of stem elongation (*t*_PH_start__ and *t*_PH_stop__) to these non-steady growth phases led to visually coherent results (Supplementary materials, Figure S1a and b, vertical lines). Nevertheless, extracting the timing of jointing was only possible for the years 2017 and 2018 when early measurements before jointing were available. These early measurements are essential for the QMER method to determine the time point when the growth rate first exceeds a certain threshold. In contrast, the end of stem elongation and final height could be extracted for all years, as late measurements after the stop of stem elongation were available.

Repeatabilities for year specific adjusted genotype means were close to 1.0 for final height, and above 0.68 for the timing of jointing and the end of stem elongation (Table 1). The heritability of across-year adjusted genotype means was highest for PH_max_ (0.98), followed by *t*_PH_stop__ (0.87) and *t*_PH_start__ (0.77). The genomic prediction accuracy for final height was with 0.78 superior to all other traits. For the timing traits, the prediction accuracies were with 0.59–0.61 lower but still strong.

**Table 1:** Repeatabilities 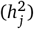, heritabilities (*h*^2^), and genomic prediction accuracies (*r*) for extracted parameters and growing seasons.

| Parameter | 2015<br>$h_j^2$ | 2016<br>$h_j^2$ | 2017<br>$h_j^2$ | 2018<br>$h_j^2$ | All<br>$h^2$ | All<br>$r$ |
| --- | --- | --- | --- | --- | --- | --- |
| $t_{PH_{\text{start}}}$ | - | - | 0.68 | 0.74 | 0.77 | 0.61 |
| $t_{PH_{\text{stop}}}$ | 0.77 | 0.86 | 0.83 | 0.71 | 0.87 | 0.59 |
| $PH_{\max}$ | 0.98 | 0.99 | 0.98 | 0.97 | 0.98 | 0.78 |
| $r_{\max}$ | 0.78 | 0.78 | 0.74 | 0.67 | 0.89 | 0.71 |
| $T_{\min}$ | 0.33 | 0.23 | 0.36 | 0.32 | 0.29 | 0.18 |
| $lrc$ | 0.57 | 0.21 | 0.52 | 0.75 | 0.63 | 0.32 |
| $lm_{\text{slope}}$ | 0.74 | 0.90 | 0.88 | 0.37 | 0.78 | 0.71 |

### 3.3. Temperature dose-response modeling allows extracting heritable curve parameters

Fitting the asymptotic model produced dose-response curves with a distinct base temperature (*T*_min_) (Supplementary materials, Figure S1c). The increase (*lrc*) in elongation rate between the base temperature and asymptote was very steep for some plots in 2016 and 2017 (e.g. plot “FPWW0120005” and “FPWW0180013” in Supplementary materials, Figure S1c) but very flat for others in 2015 and 2018 (e.g. plot “FPWW0070043” and “FPWW0220044” in Supplementary materials, Figure S1c).

For the parameter *r*_max_, indicating the maximum elongation rate, the year specific repeatabilities were generally high but lower for the years 2017 and 2018 than for other years (Table 1). High temperatures were more frequent in the years 2017 and 2018 compared to 2015 and 2016 (Figure S1d).

For the parameter *T*_min_, indicating the base temperature of growth, year specific repeatabilties were very low for the year 2016, and higher for other years with the highest value for 2017 (Table 1). Temperatures below zero were frequent for the year 2017 (with extremely low temperatures end of April) but less frequent for other years.

For the parameter *lrc*, indicating the steepness of growth response to temperature between the base temperature and maximum elongation rate, year specific repeatabilities were very low for the years 2016 but higher for other years with the highest value for 2018. The linear temperature response parameter lm_slope_ that incorporates both response to temperature and growth at optimum temperature had a very high across-year heritability. Nevertheless, it showed a large variation in repeatability between years with a very low value (0.37) for the 2018 season.

In general, repeatabilities revealed large variations for temperature dose-response curve parameters among years (Table 1). Nonetheless, the across-year heritability was above 0.63 for the two traits *r*_max_ and *lrc*, indicating a strong physiological basis. This finding was further confirmed by the strong genomic prediction accuracy of 0.71 for *r*_max_. Nevertheless, the prediction accuracy of *T*_min_ and *lrc* were lower (0.18–0.32).

### 3.4. Grouping by country of registration reveals trends of selection

Significant effects of registration country were observed for the two phenology traits, all temperature response parameters, and PH_max_ (Figure 5). The genotype means for PH_max_ (final height) and *r*_max_ (growth rate at optimum temperature) showed the same pattern of AT/CZ ≥ PL > DE ≥ CH > SE/DK ≥ FR ≥ GB, and consequently, the largest differences for PH_max_ (0.28 m) and *r*_max_ (0.20 mm/h) were found between AT/CZ and GB.

**Figure 5:**
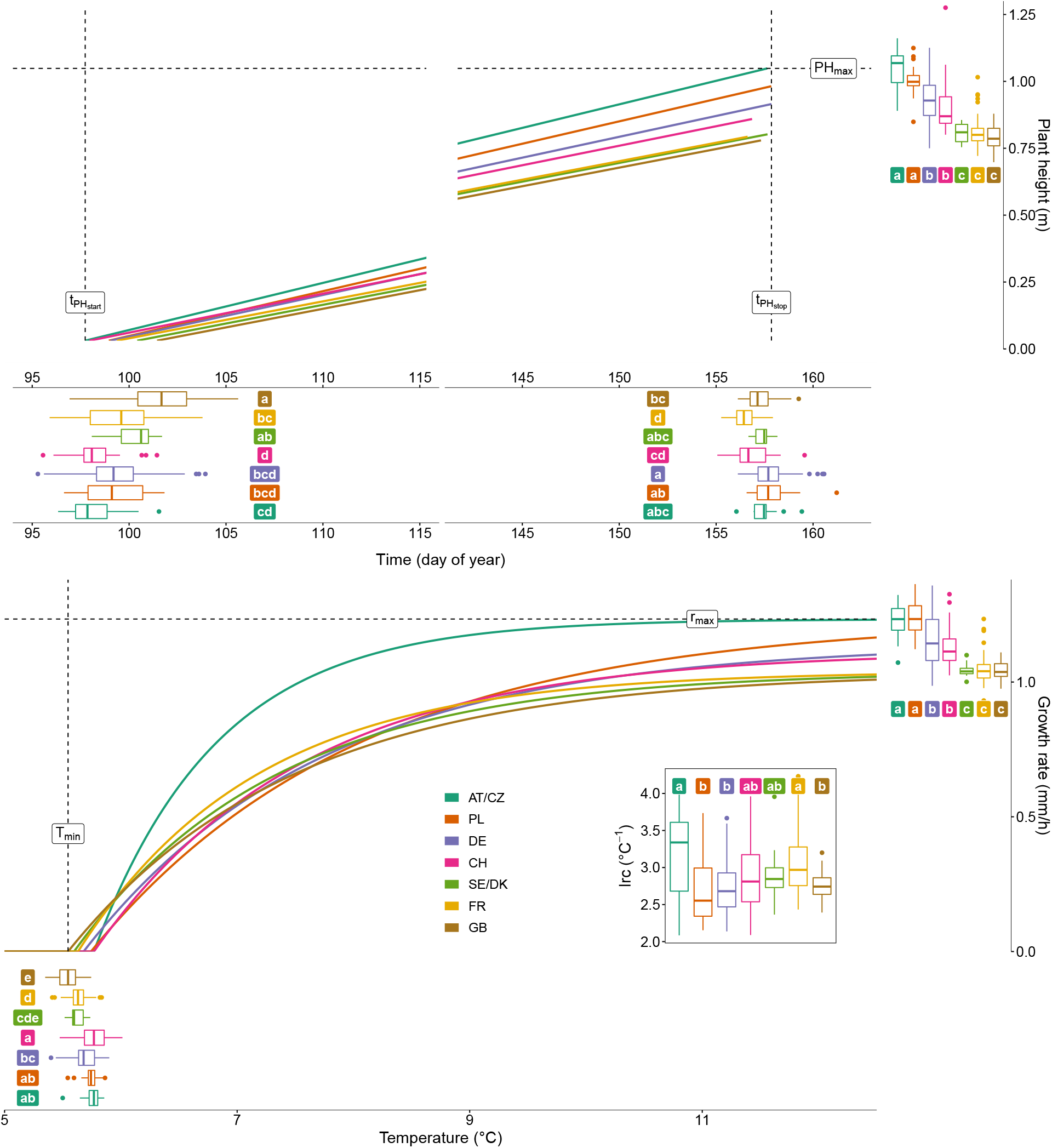
Trends of selection within country-of-registration groups for two timing traits related to jointing (*t*_PH_start__) and end of stem elongation phenology (*t*_PH_stop__) (a), for the quantity trait final height (PH_max_) (a), and for the temperature dose-response curve parameters growth at optimum temperature (*r*_max_), minimum temperature of growth (*T*_min_), and steepness of response (lrc) (b). Box plots indicate the distribution of genotypes within country groups and solid colored lines indicate the group medians. Significant (*α* = 0.05) differences between country groups are indicated by letters (a–e; country groups with the same letter are not significantly different).

In comparison, the pattern for *t*_PH_start__ (jointing) was roughly inverted with AT/CZ ≤ CH ≤ PL ≤ DE ≤ FR ≤ SE/DK ≤ GB. Again, the largest difference was found between AT/CZ and GB (3.8 days), while CH genotypes started jointing only 0.2 days later than AT/CZ.

The steepness of temperature response parameter *lrc* showed a pattern with PL and AT/CZ on the extremes, PL ≤ DE ≤ GB ≤ SE/DK ≤ CH ≤ FR ≤ AT/CZ. Consequently, AT/CZ genotypes exhibited a significantly steeper response to temperature (3.3 °C^-1^) than PL genotypes (2.6 °C^-1^). The genotype means for *t*_PH_stop__ (end of stem elongation) showed a comparable pattern: FR genotypes were the earliest and PL and DE genotypes the latest, with the largest difference (1.3 days) between FR and DE. Finally, the temperature response parameter *T*_min_ (minimum temperature of growth) was related to PH_max_ and *r*_max_ with GB genotypes among the ones with lowest *T*_min_ and AT/CZ among the ones with highest *T*_min_.

Thus, varieties registered in GB were the latest to start jointing and had the lowest minimum temperature of growth, while CH genotypes were among the earliest to start jointing and showed, together with AT/CZ varieties, the highest minimum temperature of growth.

When visualizing the temporal trends in selection in those three country groups AT/CZ, CH and GB (Figure 6), hardly any development of PH_max_, *r*_max_, and *t*_PH_stop__ was visible for genotypes registered in GB and AT/CZ. In contrast, the selection activity in CH resulted in strong and independent changes of *lrc* and *t*_PH_stop__, and closely related changes of PH_max_ and *r*_max_. Selection in AT/CZ affected *lrc* without affecting other traits. Different trends emerged when analyzing the traits *t*_PH_start__ and *T*_min_. Here, CH and AT/CZ genotypes hardly showed any development over time, but selection activities in GB have shifted *T*_min_ to lower values, also altering *t*_PH_start__ to a later timing of jointing.

**Figure 6:**
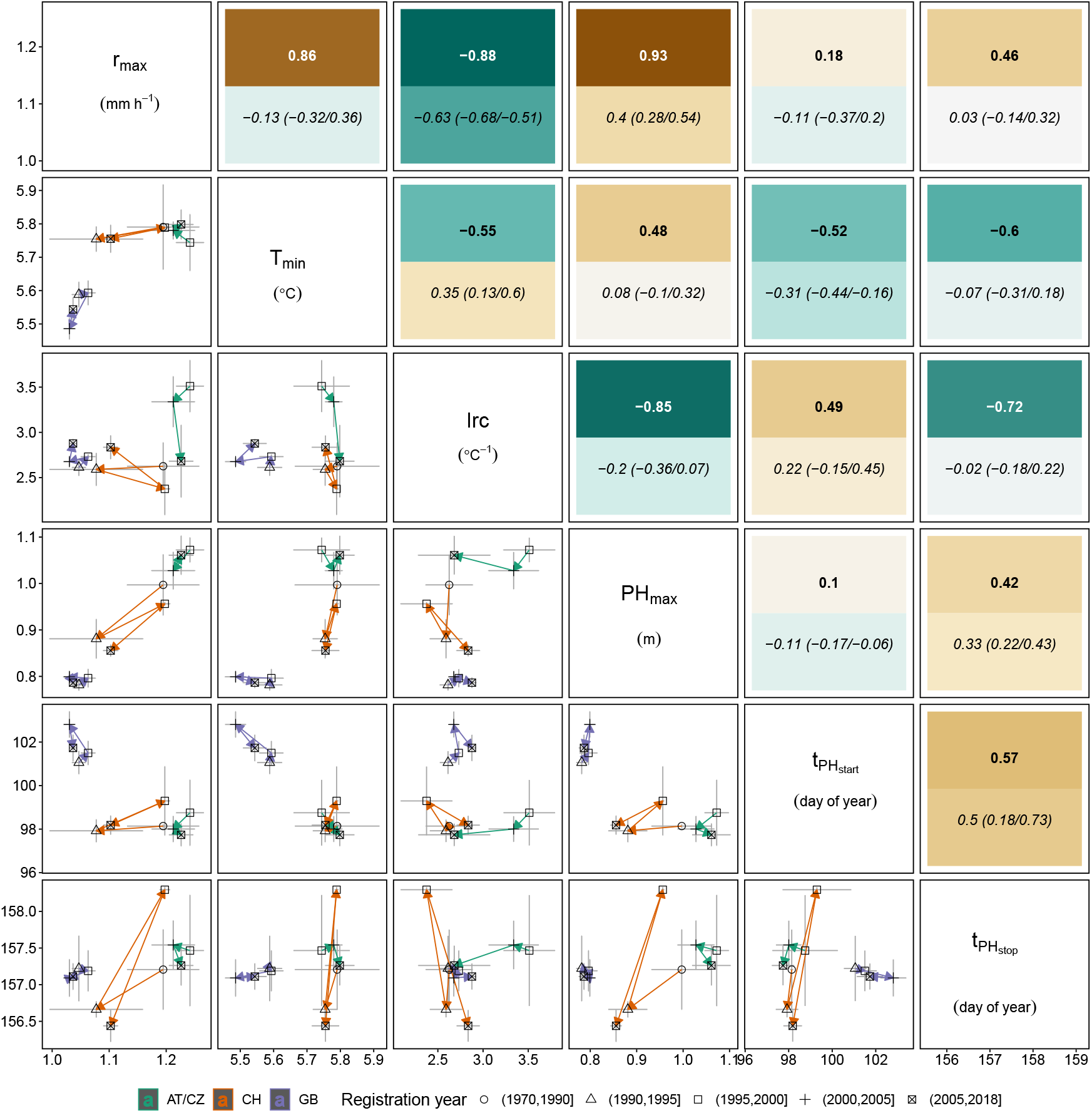
Genetic trait correlations for all 352 genotypes (upper triangle, **bold**), phenotypic correlations for all 352 genotypes (upper triangle, *italic*, mean value with minimum and maximum values indicated in brackets), and mean and standard error for genotype BLUPs of AT/CZ, CH and GB genotype (lower triangle; see color legend). The country-specific vector paths (arrows) represent the temporal development of the trait in dependence of the registration year (see symbol legend). All correlations are significant (*α* = 0.05) unless otherwise indicated (ns).

### 3.5. Trait correlations confirm connection between temperature response, phenology and height

Based on genetic correlations (Figure 6), it became evident that PH_max_ is driven by temperature response parameters (*T*_min_, *r*_max_, *lrc*) and a delayed end of stem elongation (*t*_PH_stop__), indicated by moderate to strong correlations. For the correlations between the temperature response parameters itself, *r*_max_ was strongly correlated to *T*_min_ and *lrc*. Nevertheless, there was only a moderate correlation between *T*_min_ and *lrc*, indicating that they can be selected partly independently. The end of stem elongation was moderately to strongly correlated with all three temperature response parameters and *t*_PH_start__.

In summary, a stronger growth at the temperature-optimum and a flat response to temperature delayed the end of stem elongation, which led to taller plants. A lower minimum temperature of growth delayed the end of stem elongation as well, but resulted in smaller plants. In any case, delaying the end of stem elongation has also delayed jointing.

So far, the reported correlations were based on genetic correlation calculations. Phenotypic correlations for individual years generally showed the same direction but differed in strength, with genetic correlations often being stronger (Figure 6). This finding indicates confounding year-effects that can be compensated for when screening multiple environments. Confounding effects were very evident for the correlations between the temperature response parameters and *t*_PH_stop__ for which phenotypic correlations were weak but genetic correlations strong to very strong. In two situations, phenotypic and genetic correlations were contradicting (*r*_max_ versus *T*_min_, *lrc* versus *T*_min_) with moderate to strong genetic correlations. For two other situations, phenotypic and genetic correlations were contradicting (*r*_max_ versus *t*_PH_start__, PH_max_ versus *t*_PH_start__) but genetic correlations were only weak anyway.

Thus, the results of the correlation analysis were conclusive regarding the nature of the observed relationships. To further investigate the basis of the observed correlations, we conducted univariate and multivariate GWAS to compare the traits based on marker trait associations.

### 3.6. Genome wide association studies reveal stable markers for across-year adjusted genotype means

The number of detected MTAs varied largely among traits, year-sites, and depending on the applied model (Figure 7, Supplementary materials, Figures S2-S8). No significant MTAs were detected for *T*_min_ for 2015, 2016 and the across-year BLUEs (Figure 7. Note that for *t*_PH_start__ the trait values (BLUEs) for 2015 and 2016 were missing, thus preventing the calculation of MTAs. For all traits except *t*_PH_stop__ and PH_max_, the highest number of MTAs were detected using the across-year BLUPs. Among the three applied models, the highest number of significant MTAs were detected using Blink, followed by FarmCPU. The MLM method only detected one significant MTA for the trait PH_max_ in 2017. There was considerable overlap in the detected MTAs between the three GWAS models within single year-sites and traits, as indicated by the sum of unique MTAs detected in one or more GWAS model (Figure 7, see also Supplementary Materials, Figure S1).

**Figure 7:**
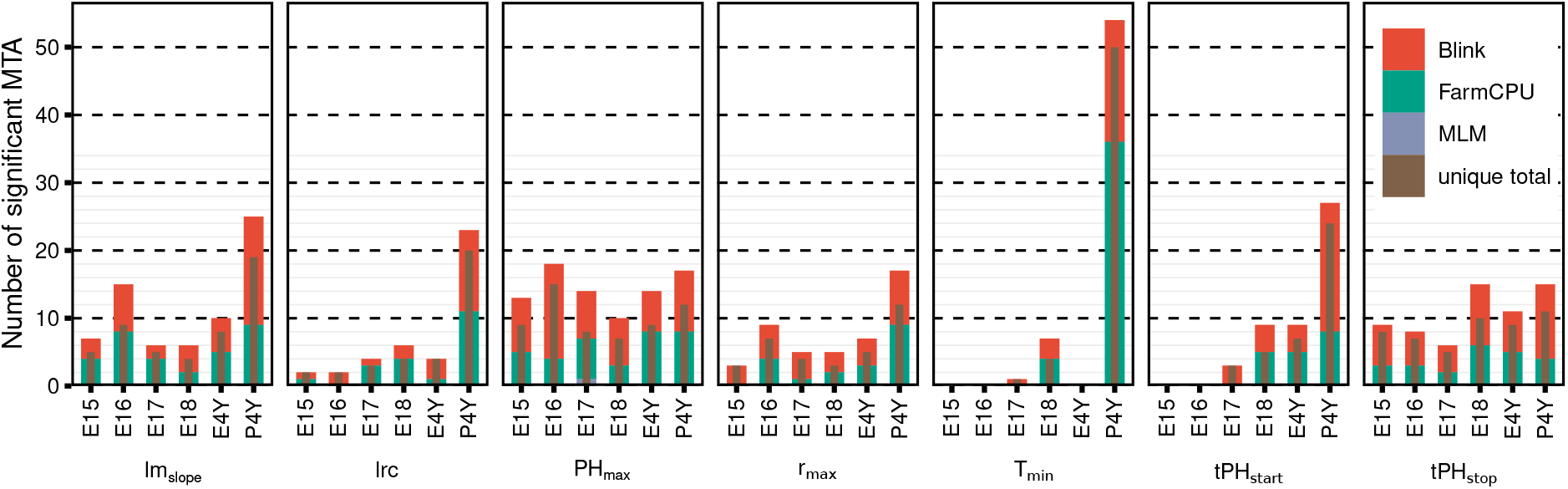
Number of detected significant marker-trait associations (MTAs) among the three univariate GWAS models (see color legend) and all traits and year genotype BLUEs (E15–E18) and across-year genotype BLUEs (E4Y) and BLUPs (P4Y). The brown bars indicate the total number of MTAs uniquely identifed within each trait-year combination among the three models.

Considering significant MTAs regardless of the GWAS model, stable markers consistently associated across several of the six analyzed models were investigated (four single year-site BLUE models, one across-year BLUE model, one across-year BLUP model) for each trait. Most stable markers were detected for PH_max_, where 11 of the 23 unique MTAs in total were detected across 2–6 analyzed models (Supplementary materials, Figures S2-S8, Table S2). For the phenology traits *t*_PH_start__ and *t*_PH_stop__, four and eight stable MTAs were detected, respectively. Among the temperature response traits, four stable MTAs were found for *r*_max_ and two for *lrc*, whereas no stable MTAs were detected for *T*_min_. The linear model for temperature response lm_slope_ yielded five stable MTAs. With the multivariate GWAS, considerably fewer significant MTAs were detected compared to the univariate GWAS and significant MTAs were only detected in the 2017 and 2018 BLUEs and across-year BLUPs (Supplementary materials, Figure S9). The multivariate GWAS on the temperature-response traits together with PH_max_ (Supplementary Figure S9, top) yielded four significant MTAs. The multivariate GWAS on the temperature-response traits together with PH_max_ (Supplementary materials, Figure S9, bottom) yielded one significant MTA.

### 3.7. Common marker-trait-associations between different traits reflect genetic correlations

To see whether the genetic correlations found in the phenotypic data are reflected in the GWAS results, all MTAs—irrespective of the year the association was detected and the GWAS model— were compared among traits. Most detected associations were unique to the respective trait. However, 17 markers in total were significantly associated in multiple univariate and multivariate GWAS models (Figure 8).

**Figure 8:**
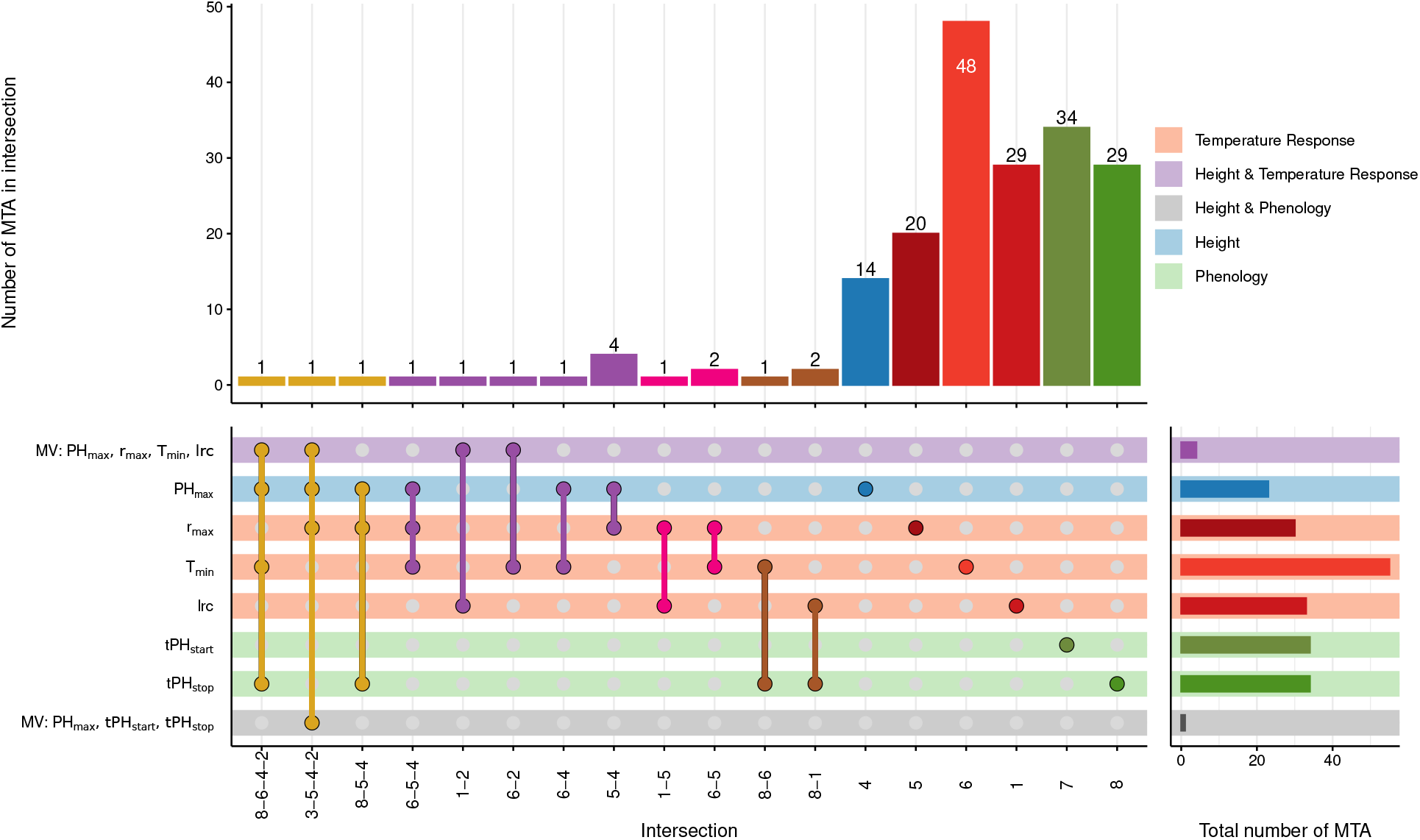
UpSet plot (Lex et al., 2014; Krassowski et al., 2021) depicting overlapping marker trait associations (MTAs) between the GWAS for the temperature-response traits *r*_max_, *T*_min_ and *lrc*, the penology traits *t*_PH_start__ and *t*_PH_stop__, PH_max_, and two multivariate GWAS combining temperature response parameters with PH_max_, and combining PH_max_ with *t*_PH_start__ and *t*_PH_stop__.

The strong genetic correlation between *r*_max_ and PH_max_ was confirmed by four common MTAs (Figure 8, intersection 5-4). Nevertheless, one common MTA for *T*_min_ and PH_max_ (intersection 6-4) and one common MTA for *T*_min_, *r*_max_ and PH_max_ (intersection 6-5-4) indicated that growth at optimum temperature was not the only driver of final height. Indeed, one common MTA each was found between the multivariate GWAS for temperature response traits / PH_max_ and the two narrow-sense temperature response parameters *T*_min_ and *lrc*, respectively (intersections 1-2 and 6-2) but no common MTA was found for *r*_max_. Although *lrc* and *T*_min_ were moderately genetically correlated, they shared no common MTA. Nevertheless, their genetic correlations to *r*_max_ were confirmed by one, respectively, two common MTAs with *r*_max_ (intersections 1-5 and 6-5).

The significant genetic correlations of *T*_min_ and *lrc* to the phenology trait *t*_PH_stop__ were confirmed by one, respectively, two common MTAs (intersections 8-6 and 8-1). The connection of *t*_PH_stop__ to the temperature response parameters but also PH_max_ was further confirmed by one common MTA among the multivariate GWAS for temperature response traits / PH_max_, *T*_min_, PH_max_ and *t*_PH_stop__ (intersection 8-6-4-2) as well as one common MTA among the multivariate GWAS for phenology traits / PH_max_, *r*_max_, PH_max_ and the multivariate GWAS for temperature response traits / PH_max_ (intersection 3-5-4-2). The common MTA between *r*_max_, PH_max_ and *t*_PH_stop__ (intersection 8-5-4) indicated that also growth at optimum temperature may influence phenology traits, independent of narrow-sense temperature response parameters.

Together, these results confirm that the investigated traits are largely independent on a genomic level. Nevertheless there are common factors between height and temperature response, temperature response and phenology as well as factors shared among all three trait groups, reflecting the pattern found in the correlation analysis.

Analysing and discussing underlying genes for the detected MTA in detail is beyond the scope of this work. Nevertheless, we searched the IWGSC refseq1.0 (Appels et al., 2018) functional annotation within chromosome specific LD windows around each MTA (supplementary file refseq_findings.xlsx, an overview is given in Supplementary Figure S10). To briefly name the most prominent genes: We detected Rht-B1 in the vicinity of Tdurum_contig64772_417 (*T*_min_-*r*_max_-PH_max_ intersection 6-5-4, distance 4.3 MB), Tdurum_contig33737_157 (*T*_min_ MTA, distance 6.8 MB) and RAC875_rep_c105718_672 (*r*_max_-PH_max_-MV(i)-MV(ii) intersection 3-5-4-2, distance 7.4 MB). Rht-D1 was found 6 MB upstream of PH_max_ MTA Kukri_rep_c68594_530 and Ppd-D1 was found 4.1 MB upstream of *lrc* MTA Excalibur_c20196_503. Furthermore, we detected Vrn-A1 6.3 MB upstream of *lrc*-QTL wsnp_Ra_c12183_19587379. These genes were mapped to the IWGSC refseq1.0 using blastn. Around the remaining MTA we detected an increased presence of genes associated with growth (i.e., gene motifs related to auxin and gibberellin (DELLA/GAI) signal transduction pathways, as well as motifs related to GRAS/S-CARECROW, WALLS ARE THIN 1, COBRA, CLAVATA3/ESR and WUSCHEL); phenology (i.e., gene motifs related to FLOWERING LOCUS T (FT), FLOWERING LOCUS C (FLC), CONSTANS (CO), FRIGIDA (FRI), AGAMOUS (AG), VERNALIZATION INSENSITIVE 3 (VIN3), VERNALIZATION 2 (At_VRN2) EMBRYONIC FLOWER 1 (EMF1), Flowering-promoting factor 1-like protein 1 (FLP1), LIGHT-DEPENDENT SHORT HYPOCOTYLS (LSH) and LAFY (LFY)); temperature response (i.e., cold response and low temperature and salt response associated protenins); and motifs associated with the circadian clock (i.e., response regulators, SENSITIVITY TO RED LIGHT REDUCED 1 (SRR1),TOC and TIC). However, no clear pattern emerged as to the trait groups of the respective MTA and the gene motifs found nearby (see Supplementary Figure S10).

## 4. Discussion

### 4.1. Field based phenotyping allows extracting robust basic physiological traits

In a previous study it was shown that frequent and accurate canopy height measurements enable the extraction of phenological stages as well as temperature response parameters (Kronenberg et al., 2020a). The latter were extracted using linear regressions between growth rates and average temperatures in the measurement interval. Within the observed data, the extracted temperature response parameters were highly heritable and allowed an accurate prediction of final height (Kronenberg et al., 2020a). However, as the model did not account for the non-linearity of the temperature response, the interpretability of the parameters was limited. In addition, Kronenberg et al. (2020a) used averaged temperatures, disregarding temperature fluctuations during measurement intervals. Considering the diurnal temperature pattern is of particular significance: Height measurements are usually done every few days. Hence, the measured growth between time points is the result of multiple diurnal covariate cycles, e.g., temperature courses. Aggregating these temperature courses to the frequency of canopy height measurements shrinks the observed temperature distribution towards the mean (Roth et al., 2022b). In soybean, diurnal temperature patterns have been shown to strongly affect leaf growth as well as carbohydrate metabolism and gene expression (Kronenberg et al., 2020b). Based on simulations, Roth et al. (2022b) demonstrated that using an asymptotic dose-response model and optimizing the parameters based on temperature courses instead of mean temperatures allows to more accurately describe temperature response in the stem elongation phase of winter wheat.

The results of this study demonstrate the applicability of such an asymptotic model for field-derived data. Both fixed platform and UAS-based canopy height measurements were equally suited for this purpose. While repeatabilities varied depending on the year, across-year heritabilities were high. Apparently, frequent very high temperatures decreased the repeatability of *r*_max_, frequent very low temperatures decreased the repeatability of *T*_min_, and varying temperature courses over the season decreased the repeatability of *lrc*. None of the seasons showed all these characteristics at the same time, but none was completely free of them either.

Consequently, depending on the temperature constellation in the respective environment, temperature response parameters are difficult to quantify with high precision. However, high across-year heritabilities and high number of significant MTAs for across-year BLUPs indicated that genotype-by-year interactions are small compared to genotype effects. Hence, temperature dose-response parameters represent robust basic physiological traits if monitored in multi-year trials. Roth et al. (2022a) could show that such basic physiological traits allow the phenomic prediction of yield, yield stability and protein content for a Swiss elite winter wheat set. The reported heritabilities and genetic correlations of this study of phenology and temperature response traits are in accordance with the ones reported in Roth et al. (2022a), raising hope that similar methods will also work on less diverse genotype sets.

### 4.2. Temperature response traits are independent drivers of phenology and height

The results revealed a clear region-driven structure within the observed population regarding the origin of the genotypes. The findings indicate that final height, phenology and temperature response traits are related to the adaptation to various climatic regions and production systems. A connection among temperature response, phenology and height was previously reported (Kronenberg et al., 2020a). While the current results confirmed these findings, using the asymptotic temperature response model further allowed dissecting and clarifying temperature response relationships and their genetic make-up.

The genetic correlations and concurring shared MTAs among *r*_max_, PH_max_, *t*_PH_start__ and *t*_PH_stop__ indicated some common genetic basis. The GWAS results further confirmed that all of these traits are highly quantitative and only a fraction of the detected QTL are shared among these traits.

It is known that the stem elongation rate of genotypes with comparable phenology but different final heights (e.g. near-isogenic lines for GA-insensitive dwarfting genes) differs (Youssefian et al., 1992). The effect of reduced height genes on growth rates was confirmed by the strong genetic correlation between *r*_max_ and PH_max_, but the correlations with other parameters and common shared MTAs indicated that dwarfing genes were not the only driver of the growth rate.

In their study investigating the effect of breeding on temperature response, Parent and Tardieu (2012) normalized the elongation rates at optimal temperatures (or better to say at 20 °C which is close to the optimum). This was done to enable the comparison of different growth processes at different scales. However, the absolute growth at optimal temperature is a relevant component of plant adaptation. In contrast to the functional temperature response model used by Parent and Tardieu (2012), the asymptotic temperature response model used in this study allowed gaining insights on the base temperature of growth, the steepness of the response, and the maximum growth rate *r*_max_.

The parameter *r*_max_ is not a temperature response in the narrow sense but rather represents the growth at the temperature optimum. In contrast, the curvature parameter *lrc* and the base temperature *T*_min_ may indicate temperature dependencies of growth. Genetic correlations as well as GWAS results indicated that *T*_min_ and *lrc* are partly independent, heritable and highly quantitative traits. Regarding the physiological basis of these traits, further research is warranted. While the reported MTAs and surrounding genome regions point towards gene motifs reported to be involved in the Arabidopsis flowering pathway (Srikanth and Schmid, 2011), these results remain inconclusive.

Based on genetic correlations and shared MTAs, one can conclude that increased *T*_min_ leads to an early end of stem elongation while only marginally affecting PH_max_. Importantly, *lrc* has a strong connection to *r*_max_ but less strong connection to PH_max_ and no common MTAs, while *r*_max_ has a strong correlation to PH_max_ and shares four MTAs. Therefore, adapting *lrc* (and, according to genetic correlations also *T*_min_) seems to have little side effect on final height. Interestingly, the steepness of response *lrc* and *T*_min_ have a much stronger influence on the end of stem elongation than *r*_max_. Consequently, temperature response in the narrow sense (*T*_min_ and *lrc*) is more closely connected to phenology than growth at optimum temperatures.

As these narrow sense temperature-response parameters appear to be partly genetically independent, these traits may be of key interest for breeding. Both final height and phenology are key traits of local adaptation. Selection for specific temperature-response trait combinations may thus allow to independently adjust phenology and height, offering opportunities towards improved adaptation to specific environments. On the downside, low genomic prediction accuracies for *T*_min_ and *lrc* based on GBLUPs signalized potential difficulties in the selection process.

In the examined set of genotypes, both phenology and temperature response appear to drive local adaptation. For varieties registered in Great Britain, selection in breeding throughout the years 1970 to 2018 led to later jointing and to decreasing the minimum temperature of growth. These two features compensated each other with respect to generating varieties of comparable height that were registered throughout the years. For varieties registered in Switzerland, plant height decreased throughout the years, coinciding with an earlier end of stem elongation and a decrease of growth at optimum temperature. In Austria and the Czech Republic, final height, jointing and end of stem elongation remained similar throughout the decades, but the steepness of temperature response increased. It has to be noted, that the observed region-specific difference may not only be driven by climatic conditions (increase in temperature and overall decrease of water availability) but also by management and policy in the particular country. Yet, the data show that country-specific strategies for the development of phenology have been selected for, allowing to keep up yield potential throughout the decades under the effect of a globally changing climate. Future studies need to reveal the physiological advantages that the observed, country-specific selection strategies have brought with them.

Yet, most clearly for the case example of varieties registered in Great Britain, the advantages of such selection strategies can be made clear: Given the applicability of the concept of temperature sum, a warming climate leads to the possibility of accumulating the same biomass in a shorter period of time. Hence, the shift of jointing towards a later time of the year can be compensated. Moreover, the decreasing base temperature even increases chances to grow vividly even during days with relatively low temperature that still remain frequent also towards later times of the season (Supplementary Material, Figure S1d).

## 5. Conclusion

In this study we could demonstrate that temperature response parameters are heritable traits with a strong physiological basis. High-throughput field phenotyping allows extracting such response curves and timing parameters for jointing and the end of stem elongation. Flexible and affordable drone and RGB hardware is as suitable as stationary phenotyping platforms such as the FIP, allowing breeders to scale up phenotyping to large breeding populations.

Nevertheless, response parameters are occasionally difficult to quantify with high precision in the field, as the efficiency of high-throughput field phenotyping will depend on temperature fluctuation during stem elongation. Combining multiple years will mitigate these limitations.

Analyzing the dependencies of traits and population structures revealed that breeding indeed has affected the phenology and temperature response of the stem elongation phase of wheat. Genotypic variances in both (narrow-sense) response to temperature but also growth rates at the optimum were indicated. Final height was not only driven by the maximum growth rate at the optimum, but also by phenology and by the responsiveness to temperature between cardinal temperatures. Although not equally strong for all traits, the measured prediction accuracies promise a high potential of genomic selection approaches for temperature response and phenology traits.

## Acknowledgement

We like to thank Léo Friedli for preparing the variety registration year and country data, Hansueli Zellweger for the field management, and Norbert Kirchgessner for operating the FIP (all ETH Zürich, Switzerland). Tanks goes to Marion Röder (IPK Gatersleben) for providing the GABI panel including marker data, and Dario Fossati for providing marker data for the Swiss varieties.

## Author contribution

L.R: Conceptualization, methodology, software, formal analysis, visualization, writing—original draft. L.K: Conceptualization, methodology, software, formal analysis, visualization, writing—original draft. H.A.: Conceptualization, methodology, funding acquisition, writing— review & editing. A.W.: Conceptualization, funding acquisition, writing— review & editing. J.H.: Methodology, software, writing— review & editing. F.E.: Conceptualization, methodology, writing—review & editing. H.-P.P.: Conceptualization, methodology, writing—review & editing. A.H.: Conceptualization, supervision, project administration, funding acquisition, writing— review & editing.

## Data availability

Unprocessed data are available from the authors upon reasonable request. Source code that support the findings of this study are openly available in the ETH gitlab repository at https://gitlab.ethz.ch/crop_phenotyping/htfp_data_processing.

