## Supplementary Materials 1 for "High-throughput field phenotyping reveals that selection in breeding has affected the phenology and temperature response of wheat in the stem elongation phase"

The following supplementary data are available online.

- Figure S1. Fitted curves to height data.
- Figure S2. Manhattan plots and quantile–quantile plots depicting the GWAS results for $r_{\text{max}}$.
- Figure S3. Manhattan plots and quantile–quantile plots depicting the GWAS results for $T_{\text{min}}$.
- Figure S4. Manhattan plots and quantile–quantile plots depicting the GWAS results for *lrc*.
- Figure S5. Manhattan plots and quantile–quantile plots depicting the GWAS results for $\text{lm}_{\text{slope}}$.
- Figure S6. Manhattan plots and quantile–quantile plots depicting the GWAS results for $\text{PH}_{\text{max}}$.
- Figure S7. Manhattan plots and quantile–quantile plots depicting the GWAS results for $t_{\text{PH}_{\text{stop}}}$.
- Figure S8. Manhattan plots and quantile–quantile plots depicting the GWAS results for $t_{\text{PH}_{\text{start}}}$.
- Figure S9. Manhattan plots and quantile–quantile plots depicting the multivariate GWAS results.
- Figure S10. Chromosome plot depicting SNP markers, marker trait associations (MTAs), and potentially interesting gene motifs.
- Table S1. Total number of significant marker trait associations (MTAs) for each trait across all years and for all traits in each year.
- Table S2. Stable marker trait associations (MTAs) over years.

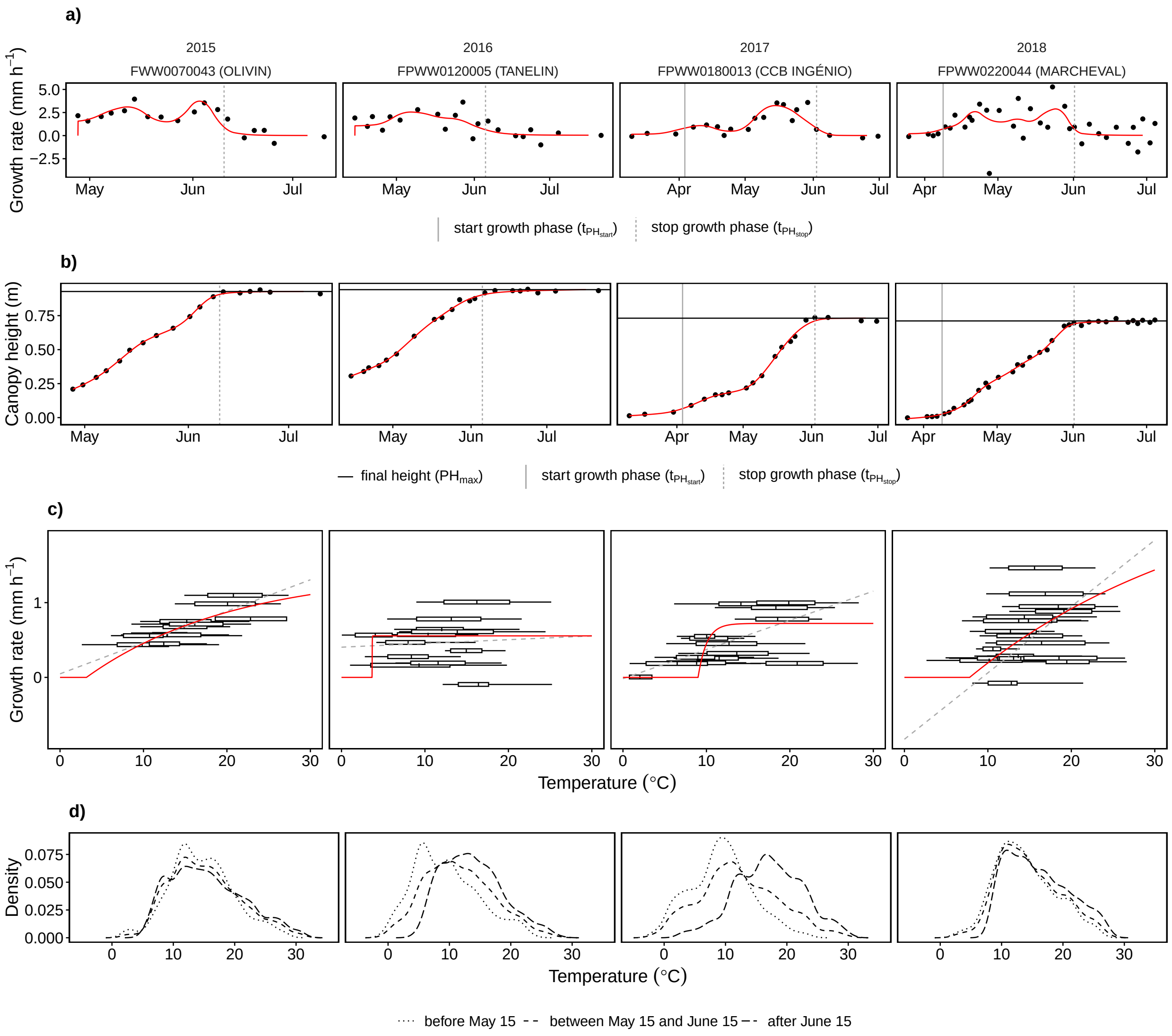


Figure S1: Growth rates (a), plant heights (b), fitted temperature dose-response curves (c), and corresponding distribution of measured temperatures (d) for four randomly selected genotypes in four growing seasons. The solid red lines indicate the fitted monotonous increasing P-spline (b), its first order derivative (a) and the fitted temperature dose-response (c), black solid points represent measurement values. In addition, the detected start and end of the growing phase (solid and dotted vertical lines) and the calculated final height (solid horizontal line) are indicated.

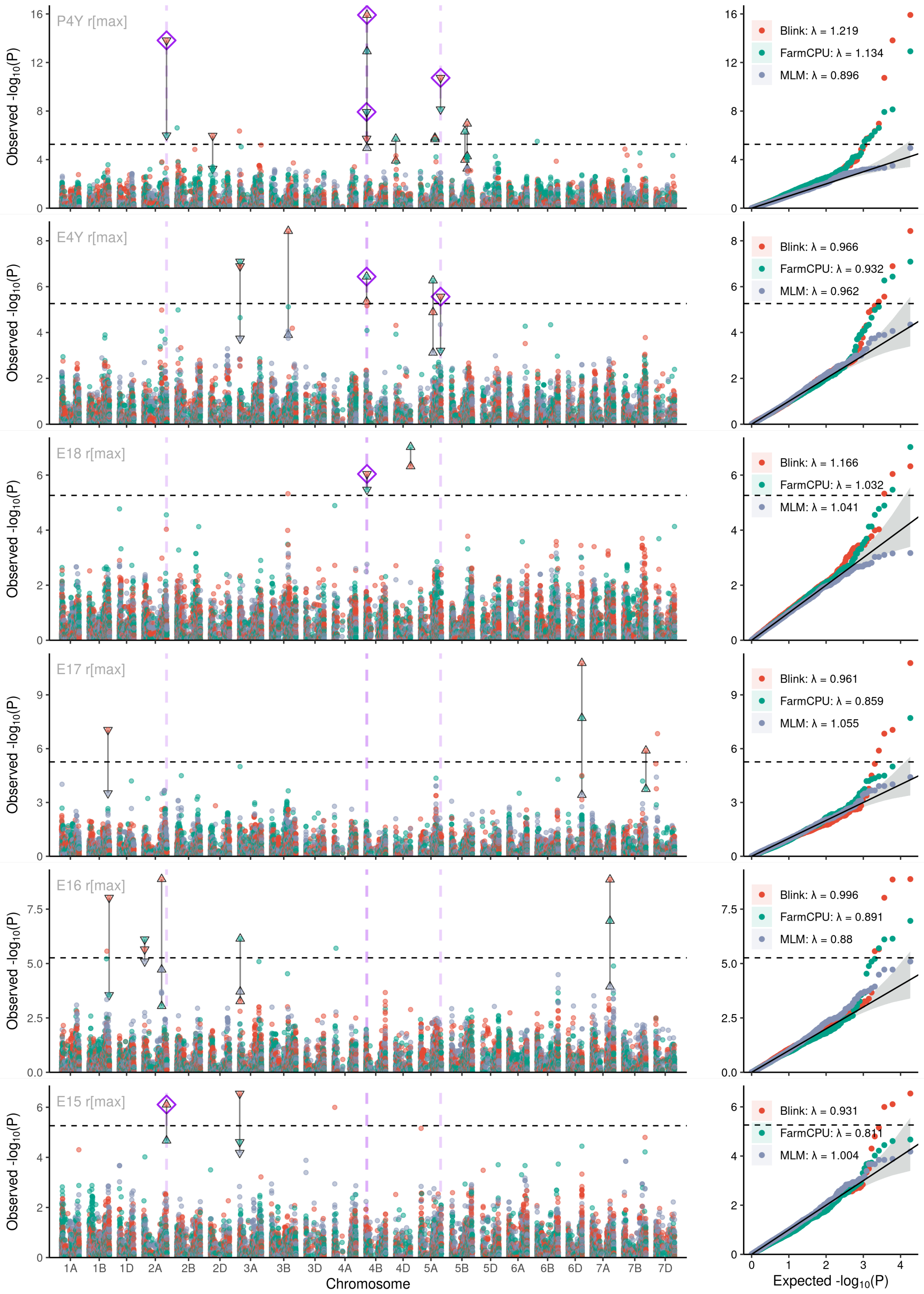


Figure S2: Manhattan plots and quantile–quantile plots depicting the GWAS results for $r_{\max}$ using the across-year BLUPs (a), across-year BLUES (b) or single-year BLUES in the respective year 2018 (c), 2017 (d), 2016 (e) and 2015 (f). In each plot, the results are given for all three applied models (Blink = red points, FarmCPU = green points, MLM = blue points) including the respective test statistic inflation factor ($\lambda$, see color legend). MTAs detected by multiple models within the respective year are indicated by black triangles connected with line segments. MTAs detected across multiple years irrespective of the GWAS model are indicated by purple rhombs and dashed vertical lines. The dashed horizontal lines indicate the Bonferroni corrected significance threshold for $\alpha = 0.05$.

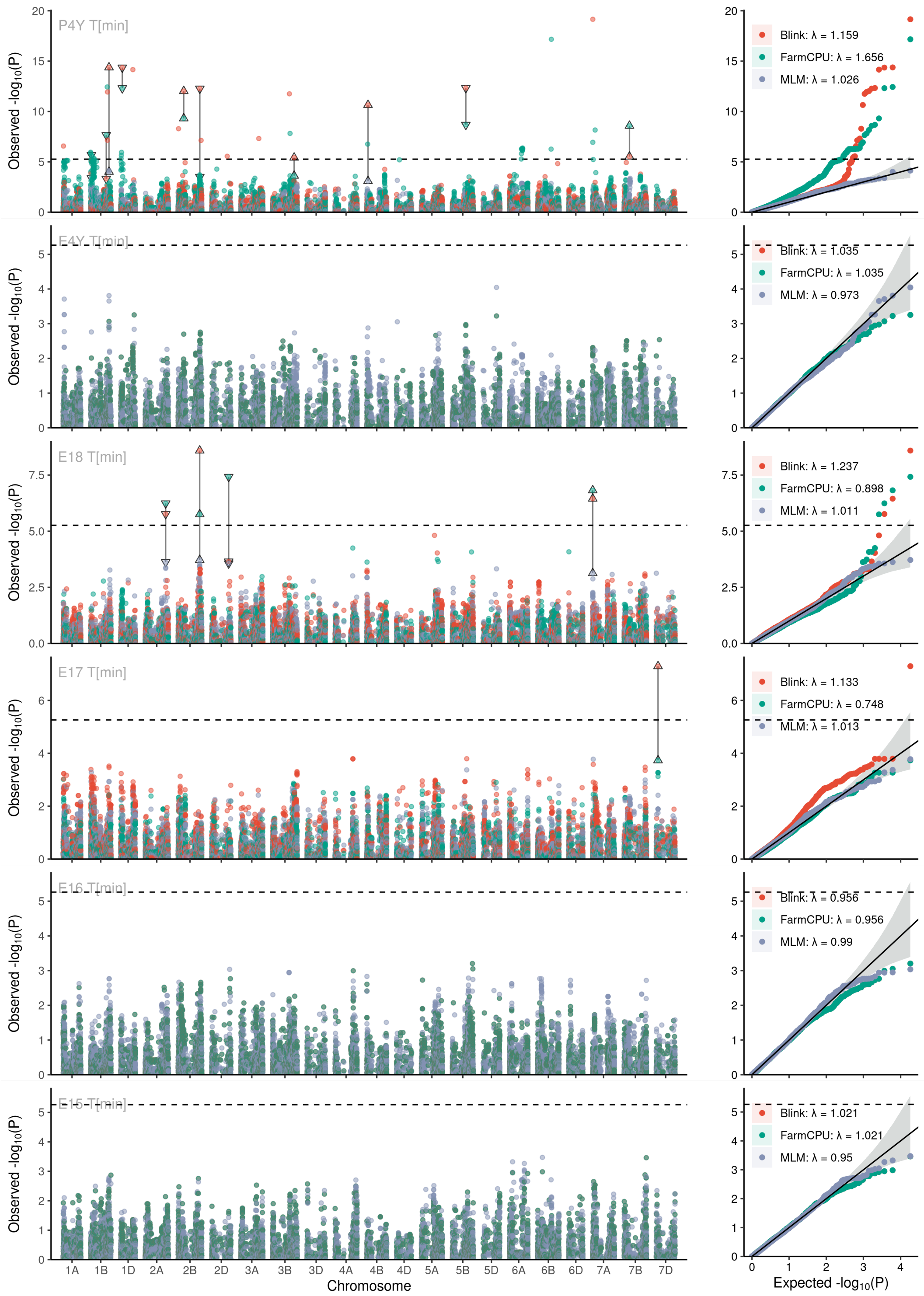


Figure S3: Manhattan plots and quantile–quantile plots depicting the GWAS results for $T_{\mathrm{min}}$ using the across-year BLUPs (a), across-year BLUES (b) or single-year BLUES in the respective year 2018 (c), 2017 (d), 2016 (e) and 2015 (f). In each plot, the results are given for all three applied models (Blink = red points, FarmCPU = green points, MLM = blue points) including the respective test statistic inflation factor ($\lambda$, see color legend). MTAs detected by multiple models within the respective year are indicated by black triangles connected with line segments. MTAs detected across multiple years irrespective of the GWAS model are indicated by purple rhombs and dashed vertical lines. The dashed horizontal lines indicate the Bonferroni corrected significance threshold for $\alpha = 0.05$. Note that the gwas resulst for Blink and FarmCPU are identical in case no significant pseudo-QTN are detected (Jiabo Wang, personal communication)

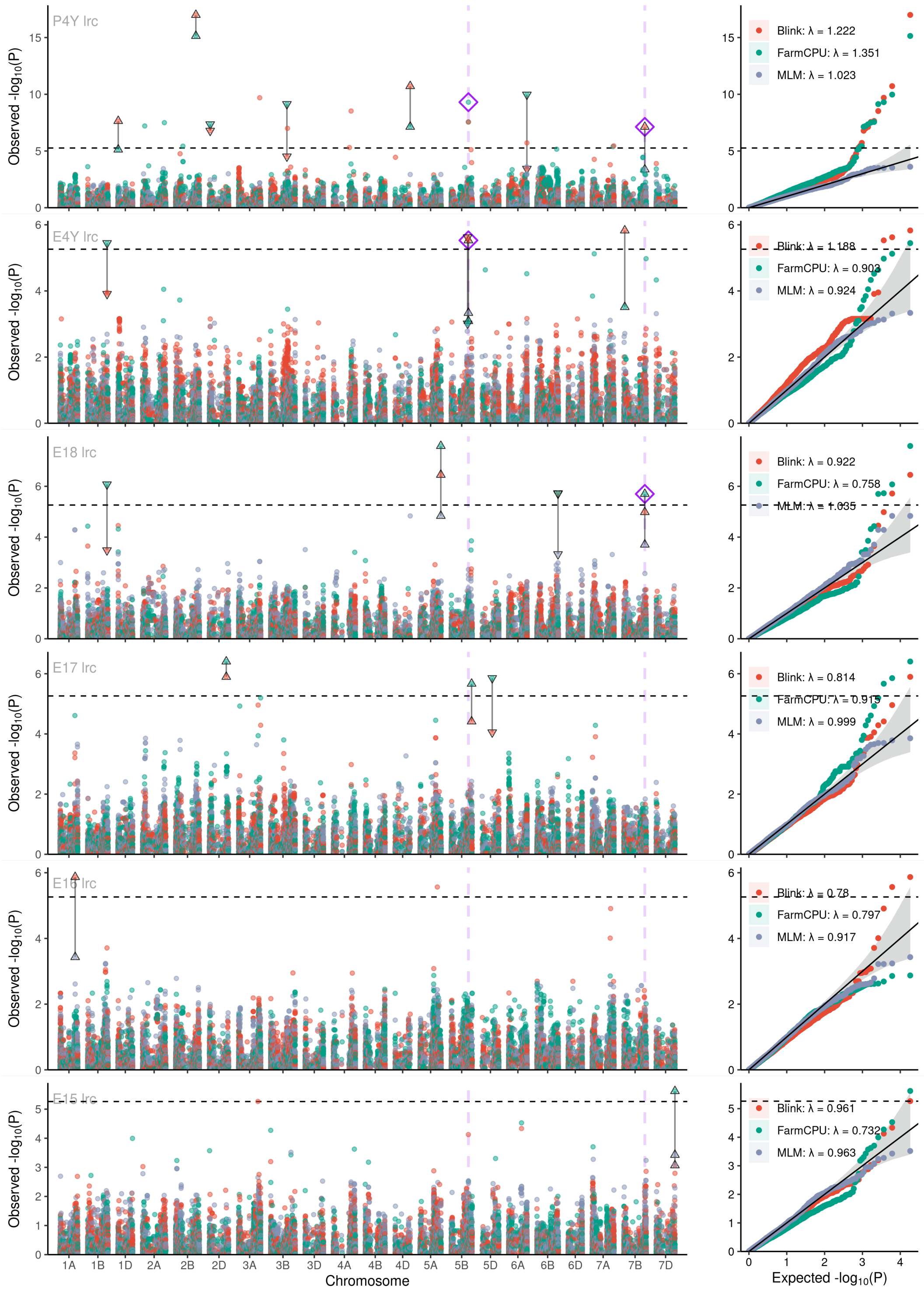


Figure S4: Manhattan plots and quantile–quantile plots depicting the GWAS results for *lrc* using the across-year BLUPs (a), across-year BLUES (b) or single-year BLUES in the respective year 2018 (c), 2017 (d), 2016 (e) and 2015 (f). In each plot, the results are given for all three applied models (Blink = red points, FarmCPU = green points, MLM = blue points) including the respective test statistic inflation factor ($\lambda$, see color legend). MTAs detected by multiple models within the respective year are indicated by black triangles connected with line segments. MTAs detected across multiple years irrespective of the GWAS model are indicated by purple rhombs and dashed vertical lines. The dashed horizontal lines indicate the Bonferroni corrected significance threshold for $\alpha = 0.05$. Note that the GWAS results for Blink and FarmCPU are identical in case no significant pseudo-QTN are detected (Jiabo Wang, personal communication)

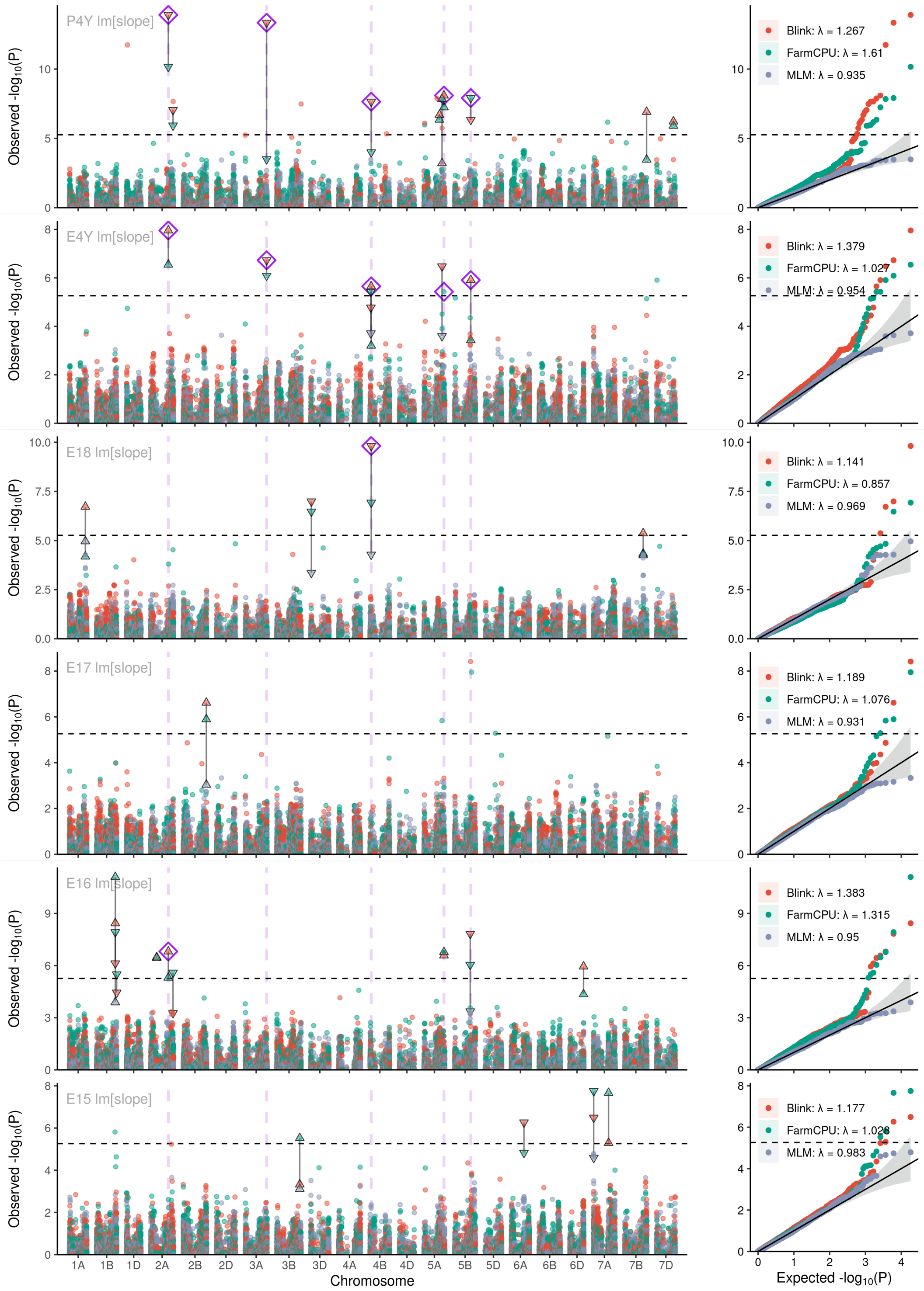


Figure S5: Manhattan plots and quantile–quantile plots depicting the GWAS results for linear temperature response model $\mathrm{lm}_{\mathrm{slope}}$ using the across-year BLUPs (a), across-year BLUES (b) or single-year BLUES in the respective year 2018 (c), 2017 (d), 2016 (e) and 2015 (f). In each plot, the results are given for all three applied models (Blink = red points, FarmCPU = green points, MLM = blue points) including the respective test statistic inflation factor ($\lambda$, see color legend). MTAs detected by multiple models within the respective year are indicated by black triangles connected with line segments. MTAs detected across multiple years irrespective of the GWAS model are indicated by purple rhombs and dashed vertical lines. The dashed horizontal lines indicate the Bonferroni corrected significance threshold for $\alpha = 0.05$.

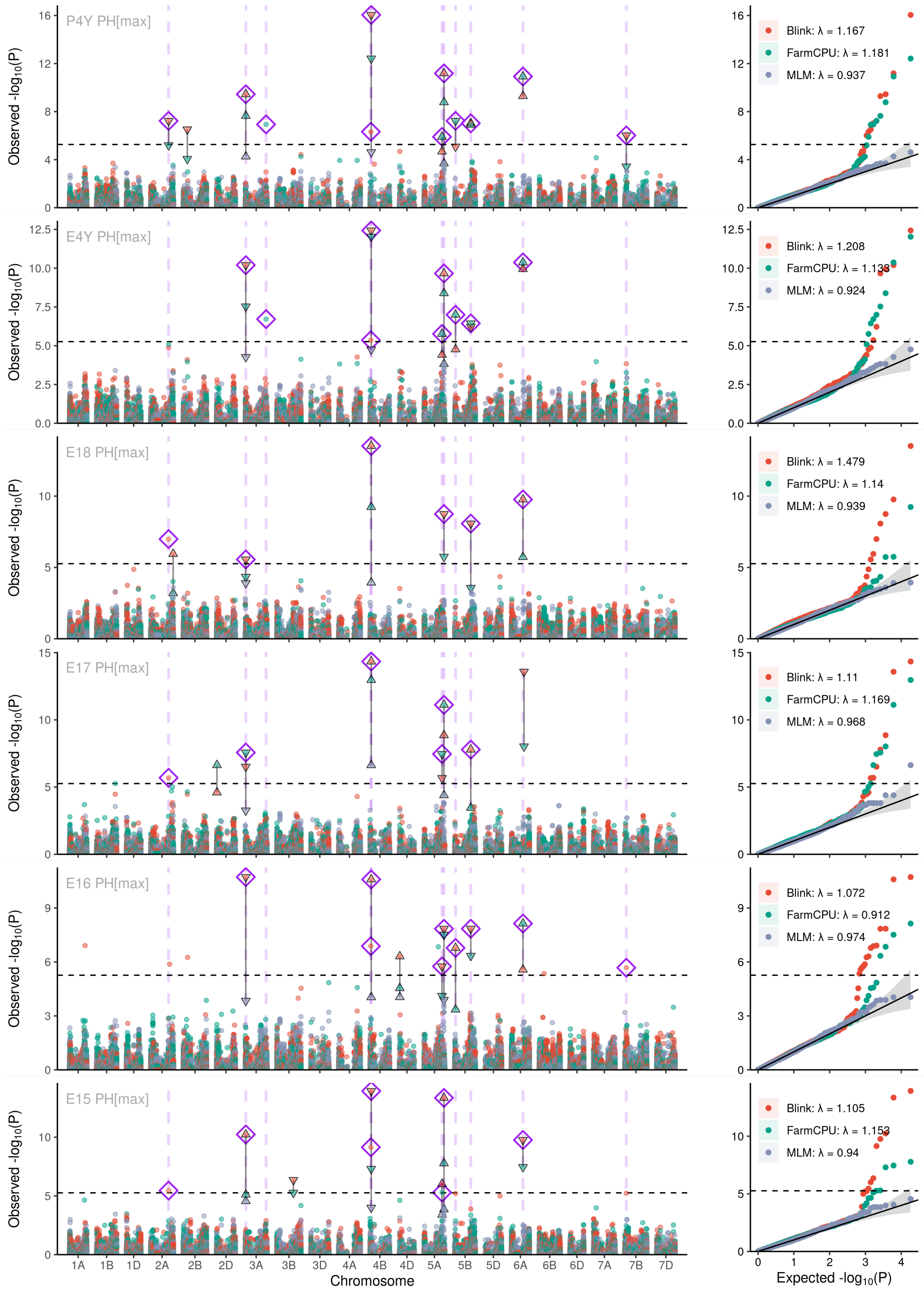


Figure S6: Manhattan plots and quantile–quantile plots depicting the GWAS results for $PH_{max}$ using the across-year BLUPs (a), across-year BLUES (b) or single-year BLUES in the respective year 2018 (c), 2017 (d), 2016 (e) and 2015 (f). In each plot, the results are given for all three applied models (Blink = red points, FarmCPU = green points, MLM = blue points) including the respective test statistic inflation factor ($\lambda$, see color legend). MTAs detected by multiple models within the respective year are indicated by black triangles connected with line segments. MTAs detected across multiple years irrespective of the GWAS model are indicated by purple rhombs and dashed vertical lines. The dashed horizontal lines indicate the Bonferroni corrected significance threshold for $\alpha = 0.05$.

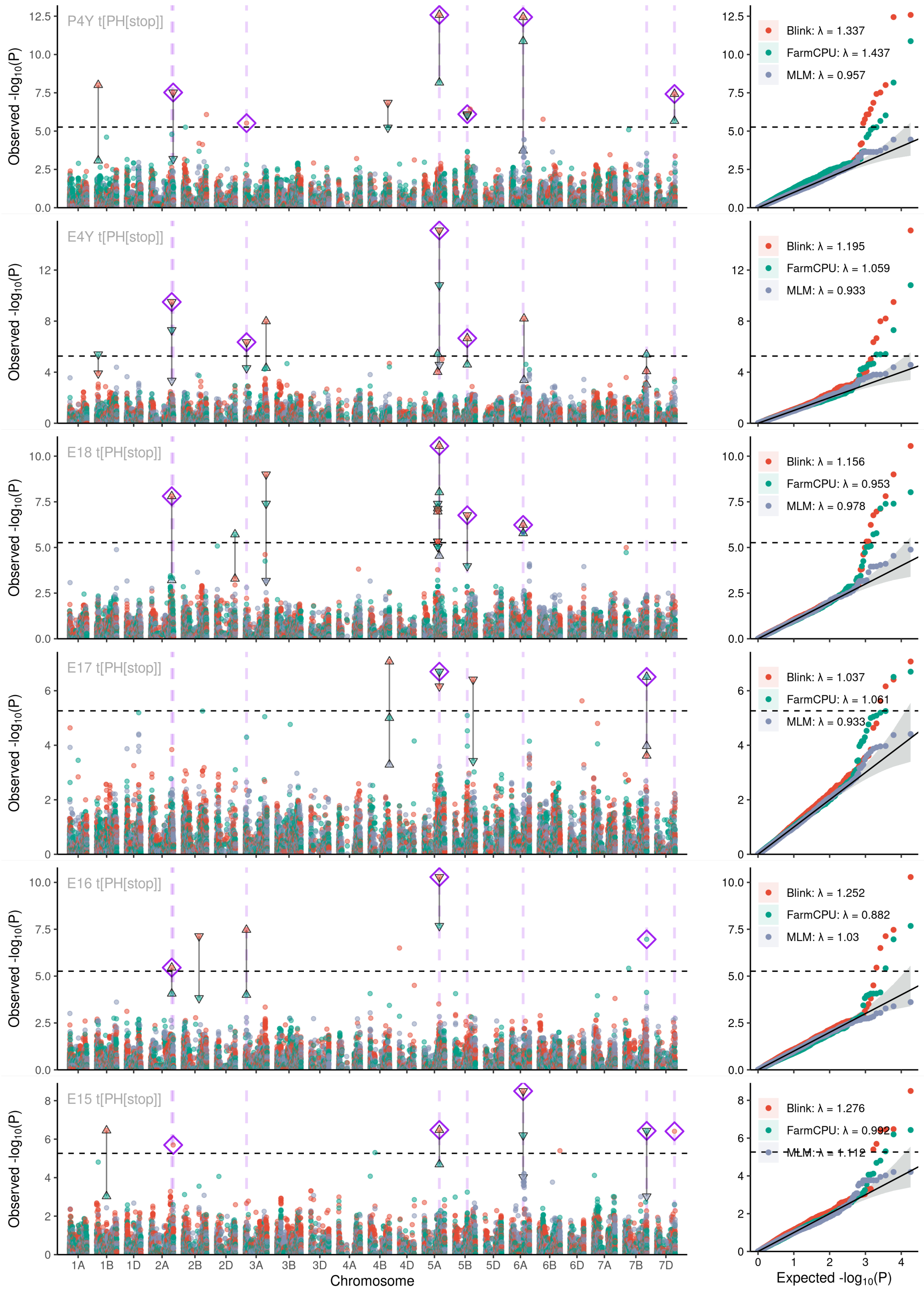


Figure S7: Manhattan plots and quantile–quantile plots depicting the GWAS results for $t_{\mathrm{PH}_{\mathrm{stop}}}$ using the across-year BLUPs (a), across-year BLUES (b) or single-year BLUES in the respective year 2018 (c), 2017 (d), 2016 (e) and 2015 (f). In each plot, the results are given for all three applied models (Blink = red points, FarmCPU = green points, MLM = blue points) including the respective test statistic inflation factor ($\lambda$, see color legend). MTAs detected by multiple models within the respective year are indicated by black triangles connected with line segments. MTAs detected across multiple years irrespective of the GWAS model are indicated by purple rhombs and dashed vertical lines. The dashed horizontal lines indicate the Bonferroni corrected significance threshold for $\alpha = 0.05$.

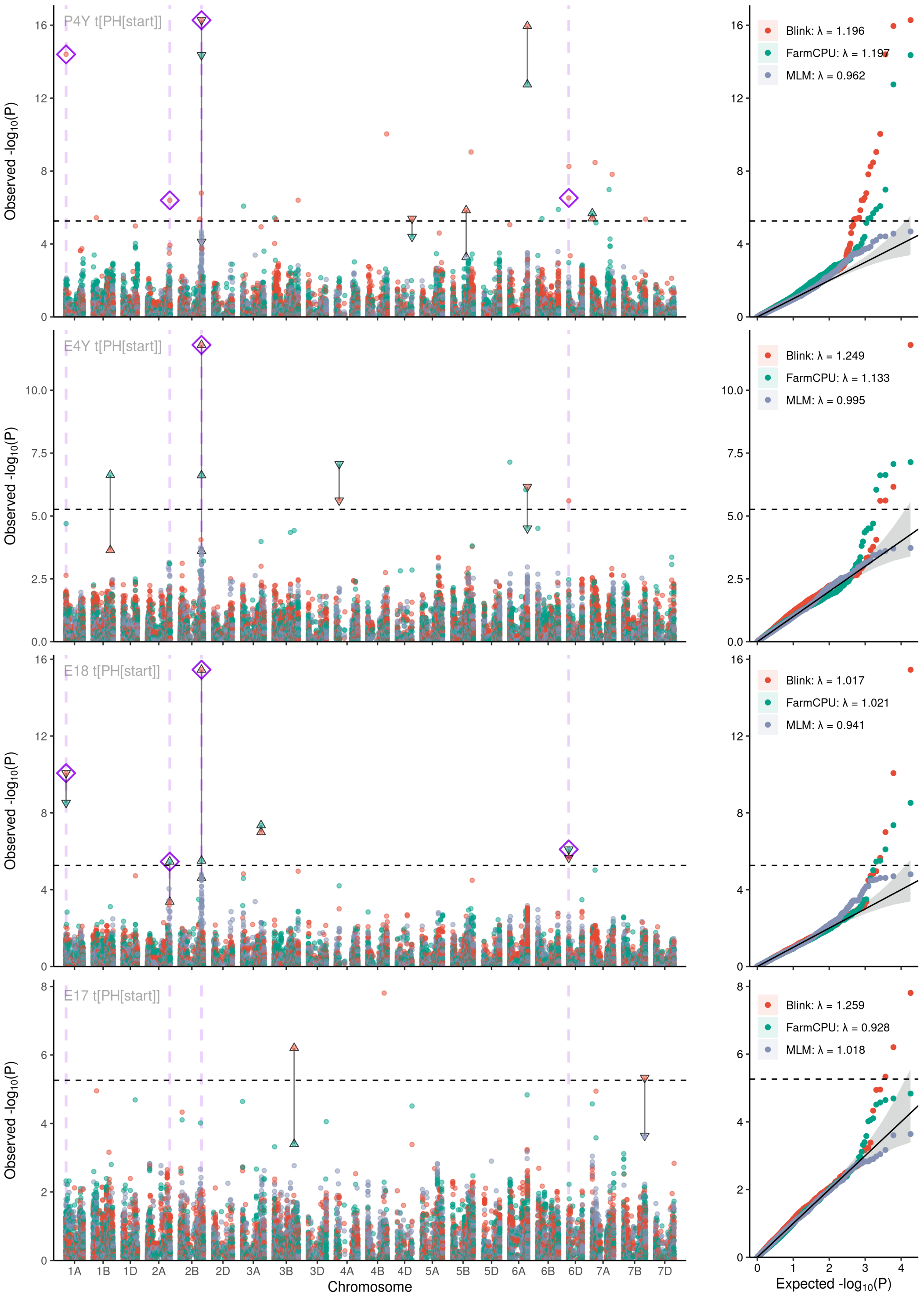


Figure S8: Manhattan plots and quantile–quantile plots depicting the GWAS results for $t_{\mathrm{PH}_{\mathrm{start}}}$ using the across-year BLUPs (a), across-year BLUES (b) or single-year BLUES in the respective year 2018 (c), 2017 (d), 2016 (e) and 2015 (f). In each plot, the results are given for all three applied models (Blink = red points, FarmCPU = green points, MLM = blue points) including the respective test statistic inflation factor ($\lambda$, see color legend). MTAs detected by multiple models within the respective year are indicated by black triangles connected with line segments. MTAs detected across multiple years irrespective of the GWAS model are indicated by purple rhombs and dashed vertical lines. The dashed horizontal lines indicate the Bonferroni corrected significance threshold for $\alpha = 0.05$.

Table S1: Total number of significant (LOD > 5.43) Marker trait associations (MTAs) for each trait across all years and for all traits in each year as well as the fraction detected by all three GWAS models (Blink, FarmCPU, MLM), two models or one model, respectively. For overlapping MTAs between the models, also non-significant MTAs (LOD > 3.00) were considered (Supplementary materials, Figures S2-S8)

| trait | yearsite | number of significant MTA | fraction detected in 3 models (%) | fraction detected in 2 models (%) | fraction detected in 1 model (%) |
|---|---|---|---|---|---|
| **all traits** | all yearsites | 323 | 18 | 42 | 40 |
| **$lm_{slope}$** | all yearsites | 50 | 20 | 48 | 32 |
| ***lrc*** | all yearsites | 35 | 14 | 43 | 43 |
| **$PH_{max}$** | all yearsites | 60 | 27 | 47 | 27 |
| **$r_{max}$** | all yearsites | 34 | 29 | 47 | 24 |
| **$T_{min}$** | all yearsites | 55 | 7 | 20 | 73 |
| **$t_{PH}start$** | all yearsites | 39 | 8 | 33 | 59 |
| **$t_{PH}stop$** | all yearsites | 50 | 18 | 58 | 24 |
| **all traits** | E15 | 27 | 30 | 33 | 37 |
| **all traits** | E16 | 40 | 20 | 48 | 32 |
| **all traits** | E17 | 29 | 24 | 48 | 28 |
| **all traits** | E18 | 37 | 43 | 51 | 5 |
| **all traits** | E4Y | 42 | 26 | 57 | 17 |
| **all traits** | P4Y | 148 | 5 | 34 | 61 |

Table S2: Number of marker trait associations detected across multiple years (including across-year BLUEs and BLUPs) for each trait.

| trait | total number of MTA | total number of unique MTA | total number of multiple yearsite MTA | average number of yearsites per MTA | fraction of multiple yearsite MTA |
|---|---|---|---|---|---|
| **$PH_{max}$** | 60 | 23 | 11 | 4 | 0.48 |
| **$t_{PH}stop$** | 50 | 34 | 8 | 3 | 0.24 |
| **$r_{max}$** | 34 | 30 | 4 | 2 | 0.13 |
| **$lm_{slope}$** | 50 | 43 | 5 | 2 | 0.12 |
| **$t_{PH}start$** | 39 | 34 | 4 | 2 | 0.12 |
| ***lrc*** | 35 | 33 | 2 | 2 | 0.06 |
| **$T_{min}$** | 55 | 55 | 0 | 1 | 0.00 |

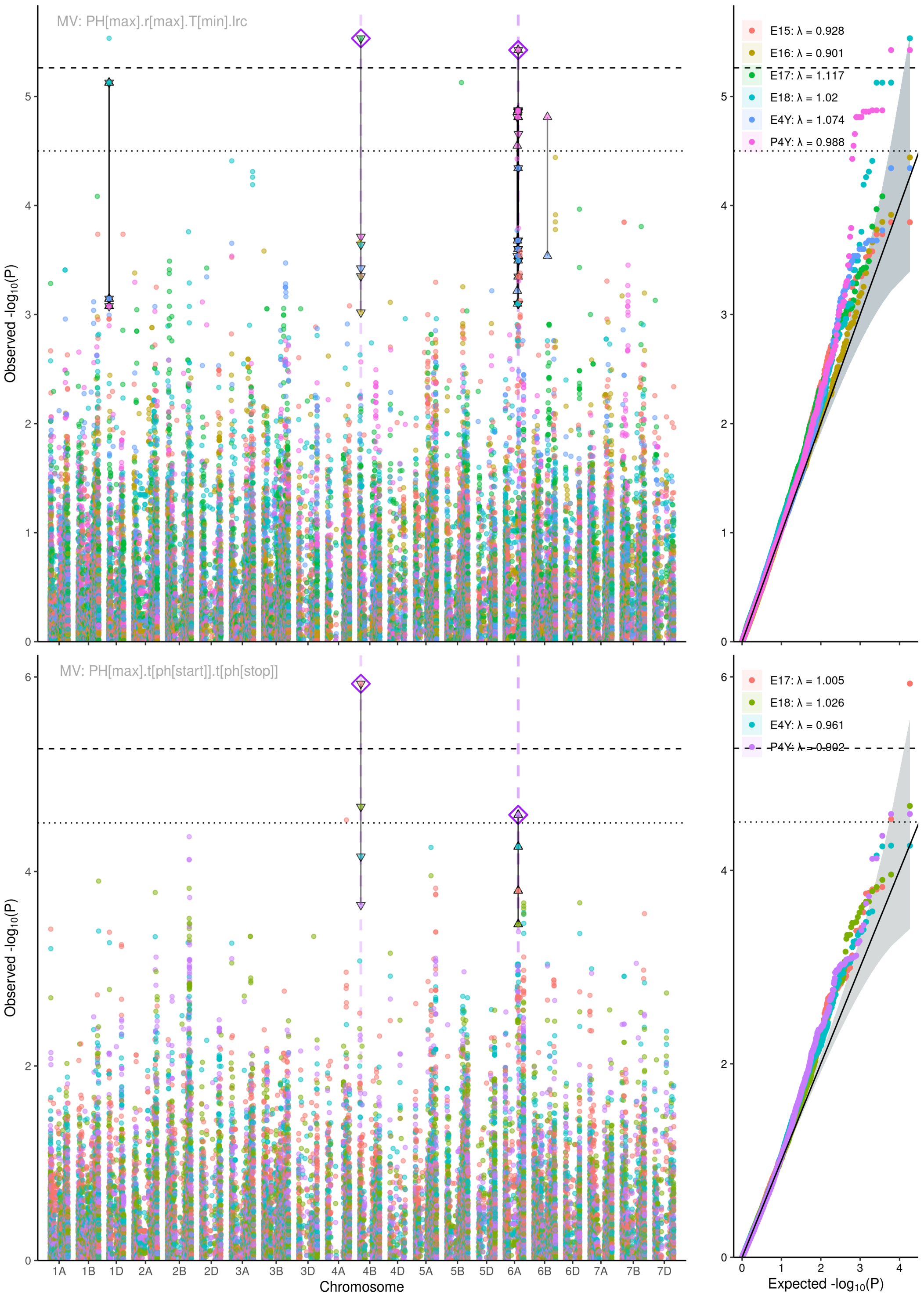


Figure S9: Manhattan plots and quantile–quantile plots depicting the multi-variate GWAS results for: (a) the temperature-response traits $r_{\max}$,$T_{\min}$,$lrc$; (b) the temperature-response traits and $PH_{\max}$, (c) all traits ($r_{\max}$,$T_{\min}$,$lrc$, $PH_{\max}$, $t_{\mathrm{PH_{stop}}}$ and $t_{\mathrm{PH_{start}}}$); (d) $PH_{\max}$ combined with the phenology traits $t_{\mathrm{PH_{stop}}}$ and $t_{\mathrm{PH_{start}}}$; and phenology traits $t_{\mathrm{PH_{stop}}}$ and $t_{\mathrm{PH_{start}}}$. In each plot, the results are given for each year (see color legend) including the respective test statistic inflation factor $\lambda$. MTAs detected across multiple models irrespective of the yearsite are indicated by purple rhombs and dashed vertical lines. These include also non-significant MTAs above the visually determined threshold of LOD > 4.5. The dashed horizontal lines indicate the Bonferroni corrected significance threshold for $\alpha = 0.05$ (i.e. LOD > 5.43).

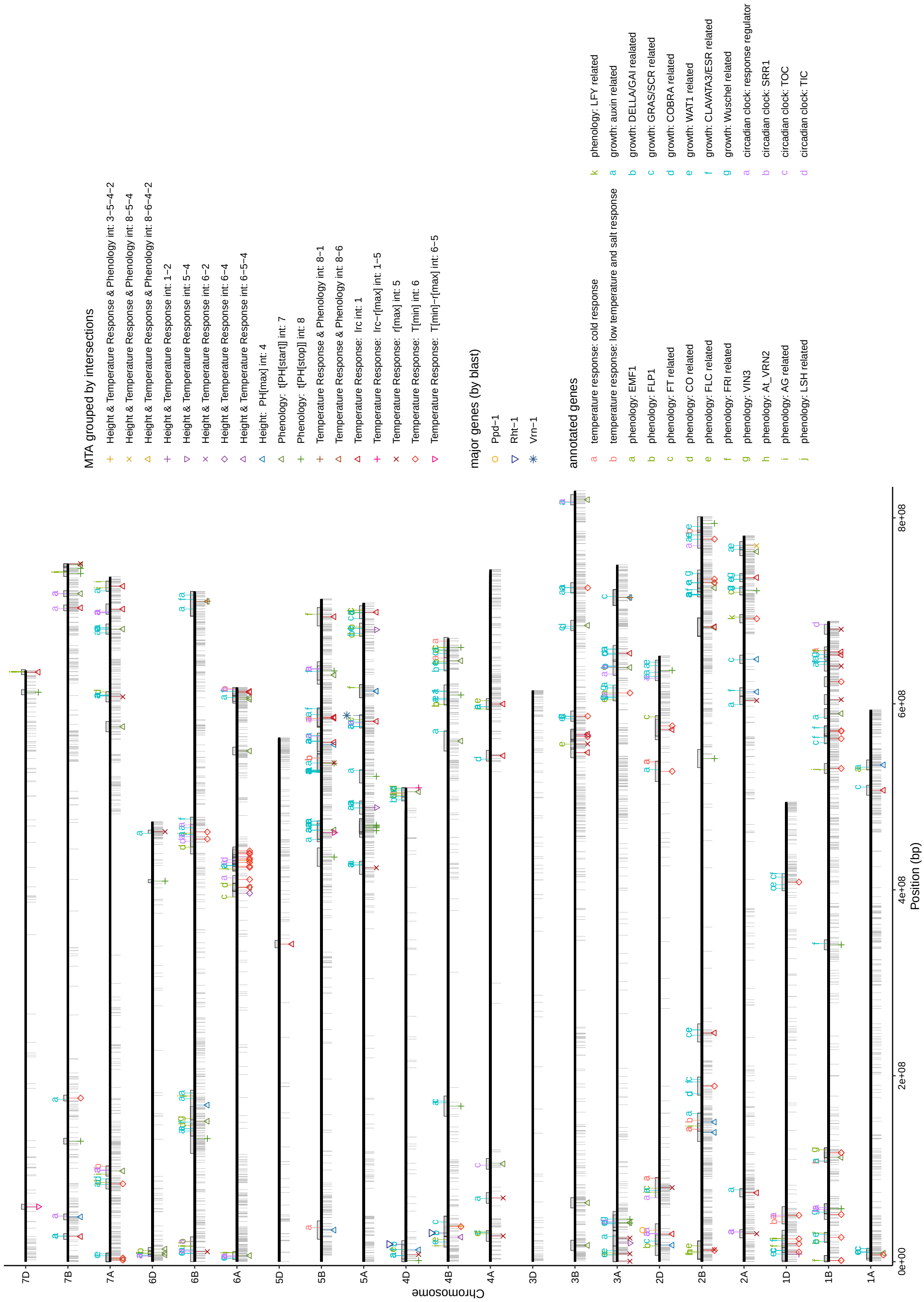


Figure S10: Chromosome plot depicting all SNP markers (grey ticks, lower side of chromosomes), detected MTA grouped by trait category intersections (lower side of chromosomes; see color legend) and potentially interesting gene motifs (upper side of chromosomes; see color legend) that were found in the IWGSC refseqv1.0 functional annotation within chromosome specific LD windows around the respective MTA (indicated by grey boxes on the upper side of the chromosomes).
